# Full-length structure of CPR-containing self-sufficient cytochrome P450

**DOI:** 10.1101/2025.11.29.689641

**Authors:** Zhenzhen Xie, Ziwei Liu, Siyu Li, Kangwei Xu, Jian-Wen Huang, Jian Min, Qiru Li, Jingxue Zhai, Te Wang, Yutong Wang, Lu Yang, Junjie Duan, Junjie Chen, Ruibo Wu, Chun-Chi Chen, Rey-Ting Guo

## Abstract

Cytochrome P450 (P450s) are heme-thiolate monooxygenases that exploit electrons sourced from pyridine nucleotide to reduce the oxygen and heme iron to catalyze hydroxylation of inactivated C-H bond^1–4^. Self-sufficient P450s that contain the substrate-binding heme-domain and the electron-donating NADPH:cytochrome P450 reductase (CPR) domain in the same polypeptide chain are highly effective and have been engineered to catalyze various challenging reactions. The high efficacy attributes to the effective electron transfer rate, but how the electrons travel among the redox centers remains elusive owing to the lack of structural information of the full-length protein. Here, we report the structure of a homologue of the most extensively studied P450BM3 from *Shimazuella soli* (*So*P450) resolved by single particle cryo-electron microscopy (cryo-EM) and X-ray crystallography. *So*P450 primarily exists as a homodimer formed via the intertwined CPR-domains. The spatial alignment of the heme-domain that is linked via an extensive loop was also determined. Notably, a class of structure that lacks one heme-domain was identified from the cryo-EM analyses, indicating that the heme-domain is mobile. We suspected that the heme-domain could move to reach the CPR-domain for electron acquisition and built a model of putative catalytic state to reveal how the electrons are relayed from NADPH to heme. These results are of fundamental importance to understand the catalytic reaction of CPR-containing self-sufficient P450s, which shall provide critical information to the engineering and applications of these enzymes.

## Main

Cytochrome P450s have attracted much attentions in organic chemistry, pharmaceutical industry and synthetic biology owing to their capability to catalyze many challenging reactions and can be engineered to accept myriad types of substrates^5–9^. The action of P450s is to catalyze the reductive scission of molecular oxygen, leading to the insertion of one atom of the dioxygen to the substrate and reduction of the other to water. The activation of the dioxygen is mediated by the highly reactive oxo iron porphyrin known as compound I, which is formed by two electrons that successively flow to the heme ^10,11^. In majority of P450s, the electrons are sourced by reduced pyridine nucleotide (NADPH or NADH) and transferred to the heme prosthetic by two types of redox partners. One is NADPH:cytochrome P450 reductase (CPR) that contains a FAD and an FMN, and the other is phthalate dioxygen reductase (PFOR) that contains an iron-sulfur cluster and an FMN.

P450s are classified as multi-component or self-sufficient systems depending on whether the heme-domain and the redox partners are located on the same gene^12,13^. For multi-component P450s, the heme-domain and the ancillary redox proteins are encoded independently such that the addition of cognate or suitable exogenous reductase is generally required^14,15^. For self-sufficient P450s, the heme-domain and the redox partner are fused in a single polypeptide thus the searching for a matching reductase is omitted. Compared with multi-component P450s, self-sufficient P450s show higher coupling efficiency and catalytic activity^16^. This superior performance of self-sufficient P450s relies on the integrity of the protein, as reconstituting individual domains of these enzymes drastically reduces the enzyme activity^17–20^. These features also make self-sufficient P450s useful frameworks for the construction of artificial chimeras to improve the catalytic activity of multi-component P450s^21–26^.

There are two types of self-sufficient P450s, one fuses with a CPR-domain and the other a PFOR-domain, which exploit electron transfer route of NADPH→FAD→FMN→heme and NAD(P)H→FMN→[2Fe-2S]→heme, respectively. In 2020, we solved the crystal structure of a PFRO-containing self-sufficient P450 termed CYP116B46^27^. This structure demonstrates that the direct electron transfer can occur between FMN and [2Fe-2S] as the distance between these two redox centers is around 8 Å. But the edge-to-edge distance between [2Fe-2S] and heme is measured to approximately 25 Å, much longer than that is demanded for efficient direct electron transfer. We thus proposed that protein residues that line the connecting tunnel between FMN and heme could serve as media for the tunneling of the long-range electron transfer and applied mutagenesis experiments to validate the role of these residues^27,28^.

The number of CPR-containing self-sufficient P450s has rapidly expanded since the first member of this superfamily termed CYP102A1 from *Bacillus megaterium* was identified (https://cyped.biocatnet.de)^29–31^. This enzyme, also known as P450BM3, is the most extensively investigated P450, which exhibits high coupling rate and turnover rate, and its heme-domain has been engineered to conduct various reactions^32–35^. A number of structures of heme-domain, FMN-domain and FAD-domain of P450BM3 and other homologous P450s have been reported^36^, but the steric organization of each domain of the full-length protein so as the mechanism that contributes to the efficient electron transfer remain elusive. It has been proposed that the FMN-domain should undergo conformational change such that the electrons from the reduced FMN could be transferred to the heme^37^. This is mainly based on the structural studies of mammalian CPRs^38,39^ and a crystal structure of partial P450BM3 fragment that contains the heme- and FMN-domain^40^. It is worth noting that 24 amino acids between the heme- and FMN-domain of P450BM3 are missing in the latter structure, complicating the determination of the domain alignment.

It has been demonstrated that P450BM3, as well as its homologs, can dimerize in solution, and the dimer configuration is essential for the enzyme to exert catalytic activity^41–45^. Biochemical and mutagenesis experiments indicate that the electrons are transferred from the FAD-domain on one polypeptide to the FMN-domain of the other polypeptide and to the heme-domain that is on the same polypeptide chain that houses the FMN-domain^44^. The dimeric configuration of P450BM3 has been revealed through negative stain electron microscopy images and single particle cryogenic electron (cryo-EM) microscopic analyses^46,47^. However, the domain alignment remains uncertain owing to low resolution. In this study, we report the cryo-EM structure and crystal structure of a P450BM3 homologous protein to elucidate the three-dimensional structure of a CPR-containing self-sufficient P450 at atomic level.

## Results and Discussion

### Full-length structure of a CPR-containing P450

Our attempt to resolve the structure of P450BM3 failed as no crystal or cryo-EM images that can be used for structure determination was obtained. This is suspected a result of high protein flexibility as mentioned in previous reports^46,47^. Since the goal is to explore the full-length structure of CPR-containing self-sufficient P450s, we examined a panel of P450BM3 homologous proteins (**Extended Data Table 1**) and eventually obtained cryo-EM images of *So*P450 from *Shimazuella soli* (60.4 % protein sequence identity to P450BM3) that can be reconstituted for structure determination (**Extended Data Fig. 1** and **Extended Data Table 2**).

A main class of the cryo-EM structure of *So*P450 that was resolved to 2.72 Å displays a dimeric configuration as the cryo-EM maps can be modeled with two heme-domains, two FMN-domains and two FAD-domains, tentatively termed as heme-A, heme-B, FAD-A, FAD-B, FMN-A and FMN-B, respectively (**Fig. 1a**). This is consistent with the size-exclusion chromatographic analyses, which indicate that *So*P450, resembling P450BM3, exists as a dimer in solution (**Supplementary Fig. 1**). The spatial location of all domains can be unambiguously determined because the electron densities of heme, FMN and FAD prosthetics were clearly identified (**Fig. 1b**). However, assigning each domain to two individual polypeptide chains is complicated as these domains that appear to intertwine to each other are linked via extensive connecting loops. The connecting regions that link heme- and FMN-domain (N456-N496) and FMN- and FAD-domain (S650-G669) are designated as CR^I^ and CR^II^, respectively (**Fig. 1a**). For CR^II^, the electron densities of the first seven amino acids are missing whereas those from D657 to G669 are sufficiently clear for the modeling (**Extended Data Fig. 2a**). Judged from the orientation of CR^II^, FMN-A and FAD-A (and FMB-B and FAD-B) are on the same polypeptide chain (**Extended Data Fig. 2b**). As such, FMN-A and FAD-B that belong to different polypeptide chains comprise a functional CPR-domain (**Extended Data Fig. 2c and 2d**), with the FMN and FAD juxtapositioned as those in other known CPR structures (see below).

**Fig. 1.**
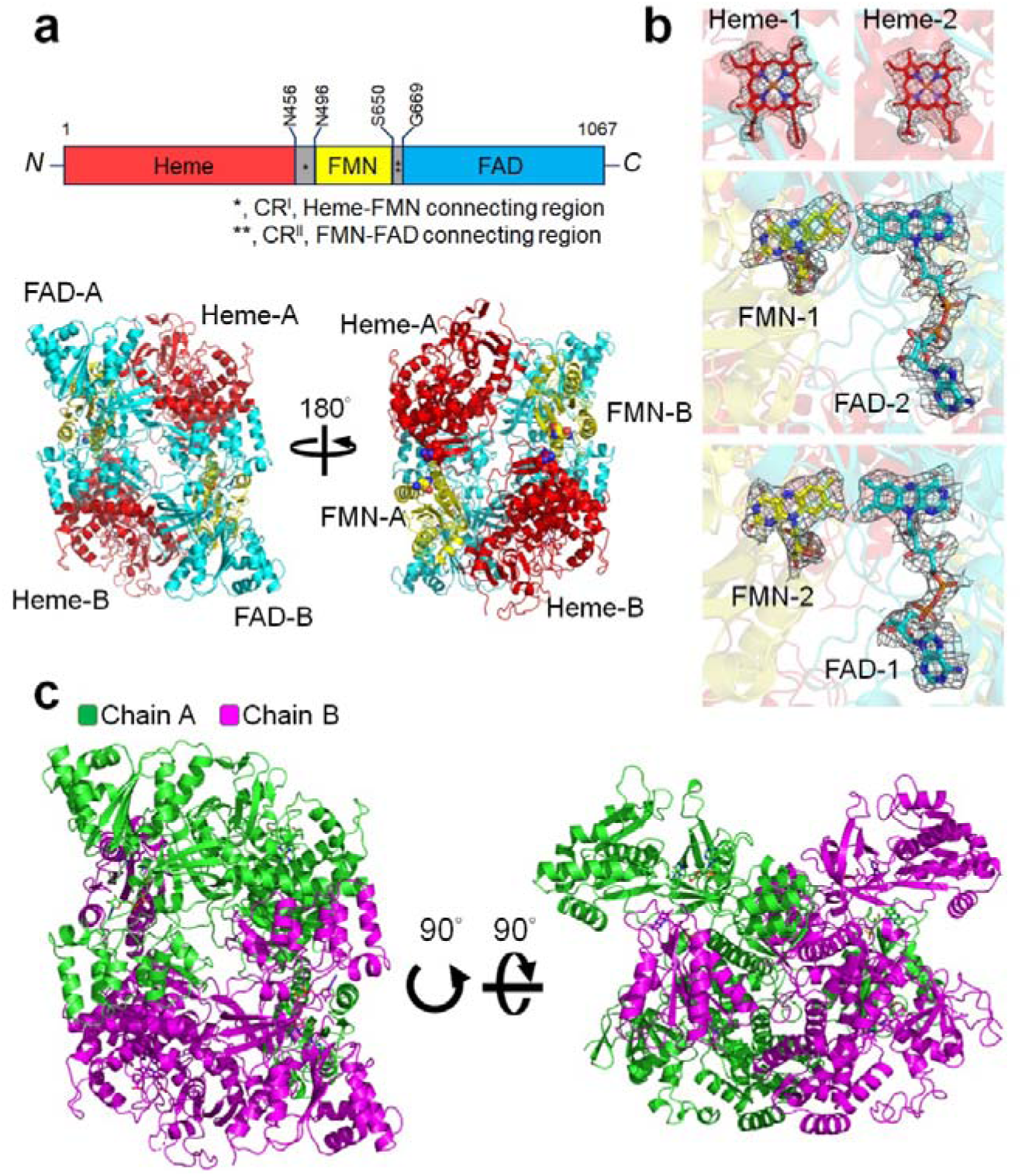
The domain assignment of cryo-EM structure of *So*P450. **a**, Top: scheme of domain distribution of *So*P450. Residues on both ends of each domain are indicated on the scheme. Linker CR^I^ and CR^II^ are indicated by asterisks. Bottom: cryo-EM structure of *So*P450 with each domain labeled and colored as the scheme shown above. **b**, The electron density maps of prosthetic groups (sticks) bound in the structure contoured at 5.0 σ are shown in mesh. **c**, Two views of *So*P450 in homodimeric configuration, with two chains colored in green and magenta.

Compared with the flavin-containing domains, the assignment of heme-domain is more challenging since the electron density of the entire CR^I^ segment is missing. Taking heme-A as an instance, it could be connected to the adjacent FMN-A via path 1 or the distal FMN-B via path 2 (**Extended Data Fig. 3a**). Despite path 1 that is shorter a more possible route, the modeling study indicates that linking heme-A to FMN-B through path 2 is also viable since CR^I^ that contains 41 amino acids (N456-N496) is sufficiently long to traverse across such a distance (**Extended Data Fig. 3b**). To resolve this issue, we constructed variant *So*P450-CR^I^-10 that contains ten amino acids in the CR^I^ region and solved its cryo-EM structure to 2.34 Å (**Extended Data Table 2** and **Extended Data Fig. 4a**). As a result, *So*P450-CR^I^-10 forms a dimer the same as the full-length protein (**Extended Data Fig. 4b**). Given that a CR^I^ that contains ten amino acids is too short to afford a dimer configured via path 2, the homodimer should be assembled through path 1 (**Supplementary Fig. 2**). As such, heme-A and FMN-A should belong to a polypeptide chain, and the structure of *So*P450 was determined (**Fig. 1c**).

In addition, we successfully grew crystals of *So*P450 and the structure was resolved at a resolution of 3.38 Å by using the template obtained in the cryo-EM analysis (**Fig. 2**). The crystal structure of *So*P450 contains two polypeptide chains that are organized as those in the cryo-EM structure (Cα root mean square deviation, 0.919 Å) (**Fig. 2a**). The electron density maps of prosthetics bound in each domain (**Fig. 2b**) and K655 to G669 and D656 to G669 in the CR^II^ of chain A and chain B, respectively, were clearly seen (**Fig. 2c**). As the case in the cryo-EM structure, no electron density map in the CR^I^ region can be seen.

**Fig. 2.**
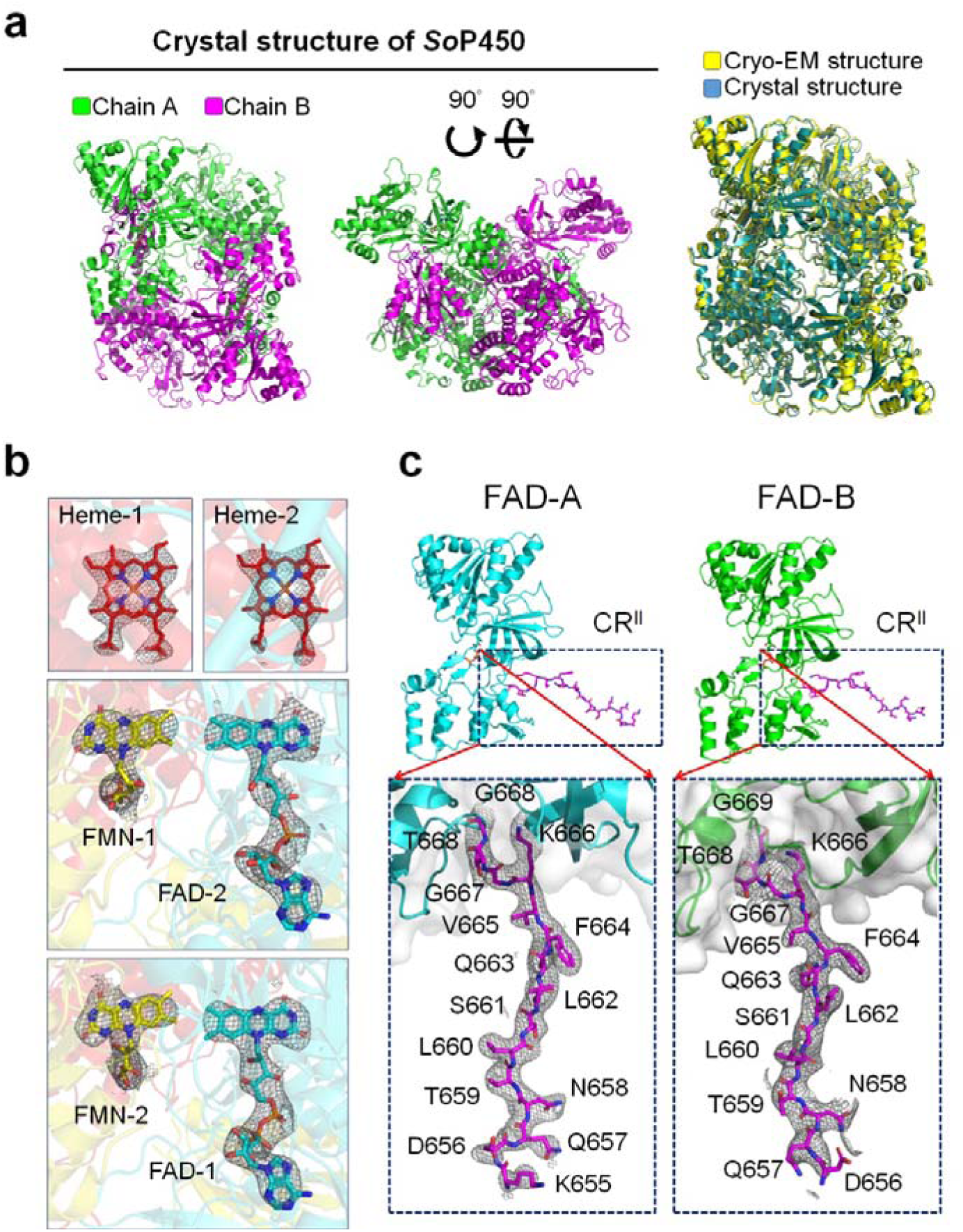
Crystal structure of *So*P450. **a**, The overall structure of crystal structure of *So*P450 and its superimposition to the cryo-EM structure of *So*P450. **b**, The 2*F*_o_-*F*_c_ electron density maps of prosthetic groups (sticks) bound in the structure contoured at 2.0 σ are shown in mesh. Each domain and the bound cofactors are displayed in a color scheme that is applied in Fig. 1c. **c**, The overall structure and 2*F*_o_-*F*_c_ electron density maps of CR^II^ segment in the crystal structure of *So*P450 contoured at 2.0 σ are shown in mesh.

### Cofactor-binding modes in *So*P450

The heme-domain located on the most N-terminus that mainly comprises α-helices adopts a canonical triangular fold of P450s (**Fig. 3a**)^12,48^. The heme is thiol-ligated to an invariant cysteine (C404 in *So*P450) on the proximal side as in other P450s^12,49^, and an elongated tunnel that should house the substrate formed on the distal side (**Fig. 3a**). The FMN-domain consists of a parallel β-sheet flanked by five α-helices with a FMN bound to the apex of the cone-shaped domain (**Fig. 3b**). The FAD-domain consists of an anti-parallel β-barrel and the putative NADPH-binding motif in another parallel five-stranded β-sheet sandwiched by several α-helices (**Fig. 3b**). We also solved the cryo-EM structure of NADPH-bound complex of *So*P450 to probe the binding mode of the nicotinamide pyridine (**Extended Data Table 2** and **Extended Data Fig. 5**). The cryo-EM structure of *So*P450/NADPH is highly identical to the apo-form structure (**Extended Data Fig. 5b**) and contains electron density maps that can be easily modeled with NADPH molecules (**Fig. 3b** and **Extended Data Fig. 5c**). The ribityl-nicotinamide moiety of the NADPH is missing, probably owing to the disordered structure of this portion as the case in other complex structures of CPR (**Fig. 3b**)^50–52^.

**Fig. 3.**
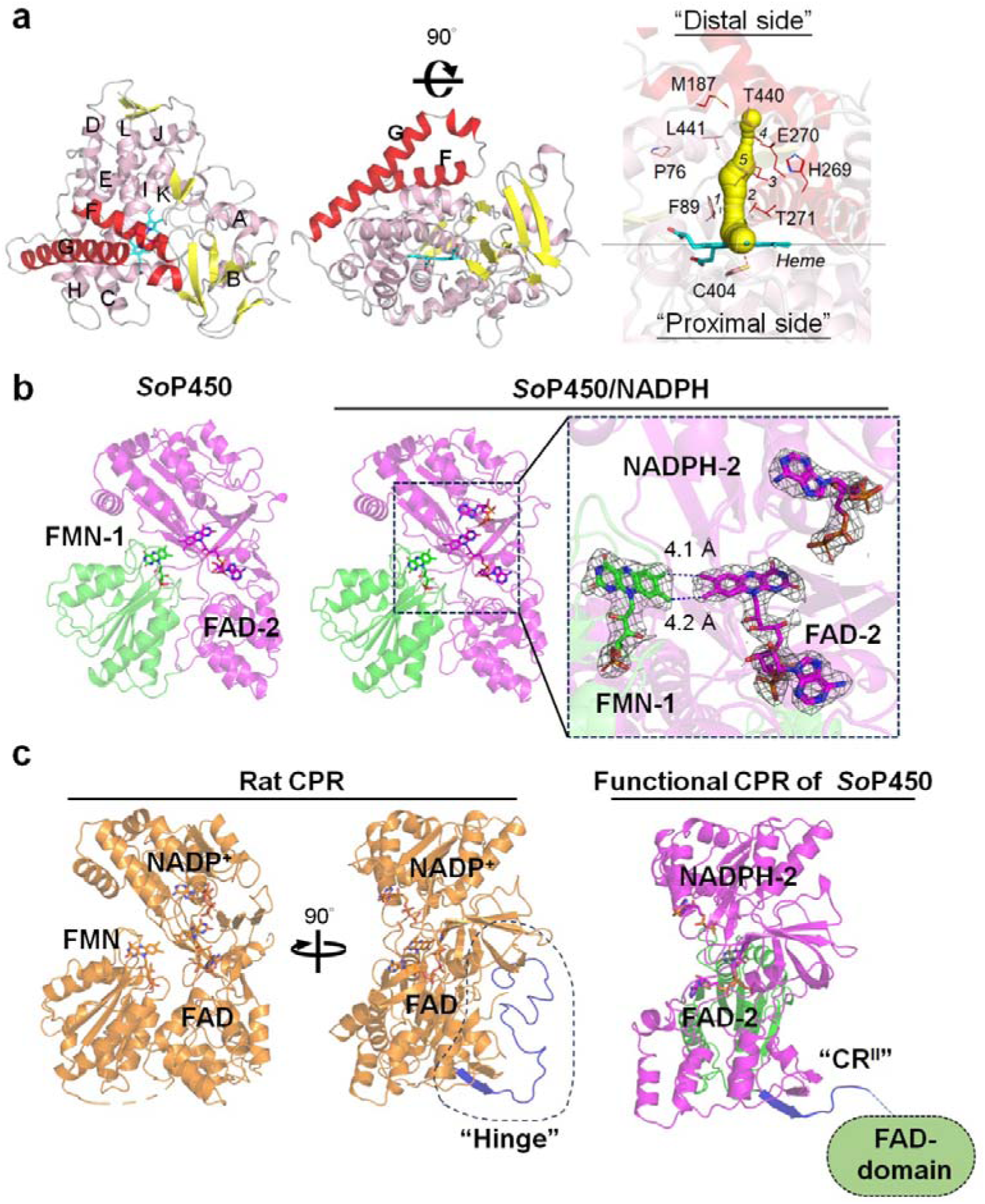
The domain structure of *So*P450. **a**, Left: two views of the heme-domain of *So*P450 with main α-helices labeled alphabetically. Right: the putative substrate-binding cavity above the prosthetic heme (cyan stick) displayed as yellow bubble. The residues lining the cavity are shown in lines. 1, A331; 2, A267; 3, I266; 4, L183; 5, T442. **b**, The di-flavin CPR domain consisting of FMN-A and FAD-B domains from different dimeric counterparts in the cryo-EM structures of *So*P450 and SoP450/NADPH are shown in green and magenta, respectively. The bound cofactors are displayed as sticks. The electron density maps of prosthetic groups bound in the complex of *So*P450/NADPH are contoured at 5.0 σ are shown in mesh and displayed in the zoom-in view. **c**, The structures of rat CPR and the functional CPR-domain of *So*P450 with hinge in the former and CR^II^ in the latter highlighted.

The FMN, FAD and NADPH bind to the protein via a number of polar interactions and the isoalloxazine and adenine are packed against several aromatic residues (**Extended Data Fig. 6a**). The complex structures of these cofactors and P450BM3 have been revealed by the crystal structures of the individual FAD-domain^42^ and a FMN-domain in complex with the heme-domain^40^, which indicate that the location and interaction networks of FAD and FMN of *So*P450 and P450BM3 are highly identical (**Extended Data Fig. 6b**). The isoalloxazine rings of two flavins are juxtaposed by their C7- and C8-methyl moieties with a distance of approximately 4 Å (**Fig. 3b**), a pose that facilitates the direct electron transfer^53^. The nicotinamide of NADPH that should pack against the isoalloxazine ring of FAD to allow the electron transfer lacks electron density maps while the indole side group of W1064 remains stacks to the isoalloxazine ring of FAD (**Fig. 3b** and **Extended Data Fig. 6a**). This suggests a competition between the nicotinamide and W1064, a phenomenon consistently seen in other complex structures of flavin-containing NAD(P)H reductase, and the nicotinamide portion could be revealed by replacing the FAD-stacking aromatic residue with small amino acids^54,55^.

The relative locations of these cofactors resemble those in CPR from other species (**Fig. 3c** and **Supplementary Fig. 3**), though the FAD- and FMN-domain in the latter group are on the same polypeptide chain^50–52^. The CR^II^ segment of *So*P450 stretches to the other CPR-domain of the dimer, whereas the equivalent region in other CPR structures forms a hinge (**Fig. 3c**). It has been suggested that the hinge region accounts for the back-and-forth movement of the FMN-domain^39,52,55^. However, such movements were not observed in the CPR-domain in all *So*P450 structures presented in this study.

### The heme-domain of *So*P450 is mobile

These structures can clearly show the electron flowing path from NADPH to FAD to FMN but that from FMN to heme would be very inefficient, if ever occurs, in the observed homodimer configuration. This is because the FMN is located distantly from either heme prosthetic with straight-line edge-to-edge distances of longer than 40 Å (**Supplementary Fig. 4**), much longer than the distance that allows efficient electron transfer^12,27^. In this context, domain rearrangement that brings FMN and heme closer is expected to take place. Structural investigations of CPRs from the multi-component P450s suggest a mobile FMN-domain that can stretch out to approach the proximal side of the heme-domain^39,56,57^. As such, FMN-A should dissociate from FAD-B, or FMN-B from FAD-A, to reach for the heme-domain. However, we did not observe movement of FMN-domain. The interaction pose of FMN-domain and heme-domain revealed in the crystal structure of partial P450BM3^40^ is also unlikely to take place in the configuration of the dimeric structure as CR^II^ segment is too short to allow the relocation of the FMN-domain (**Supplementary Fig. 5**).

Instead, we identified a class of structure that lacks one heme-domain during processing the cryo-EM images of *So*P450 (**Fig. 4a**, **Extended Data Figs. 1b and 1c**). The one-heme-missing structure, termed class 2 conformation, contains all cofactors, except for one heme, that bind to the same positions as those in the dimeric structure (**Fig. 4b**). The class 2 conformation superimposes well to the dimer though FAD-B slightly deviates from the position in the dimeric form, which could owe to the removal of the heme-A domain located beneath (**Supplementary Fig. 6**).

**Fig. 4.**
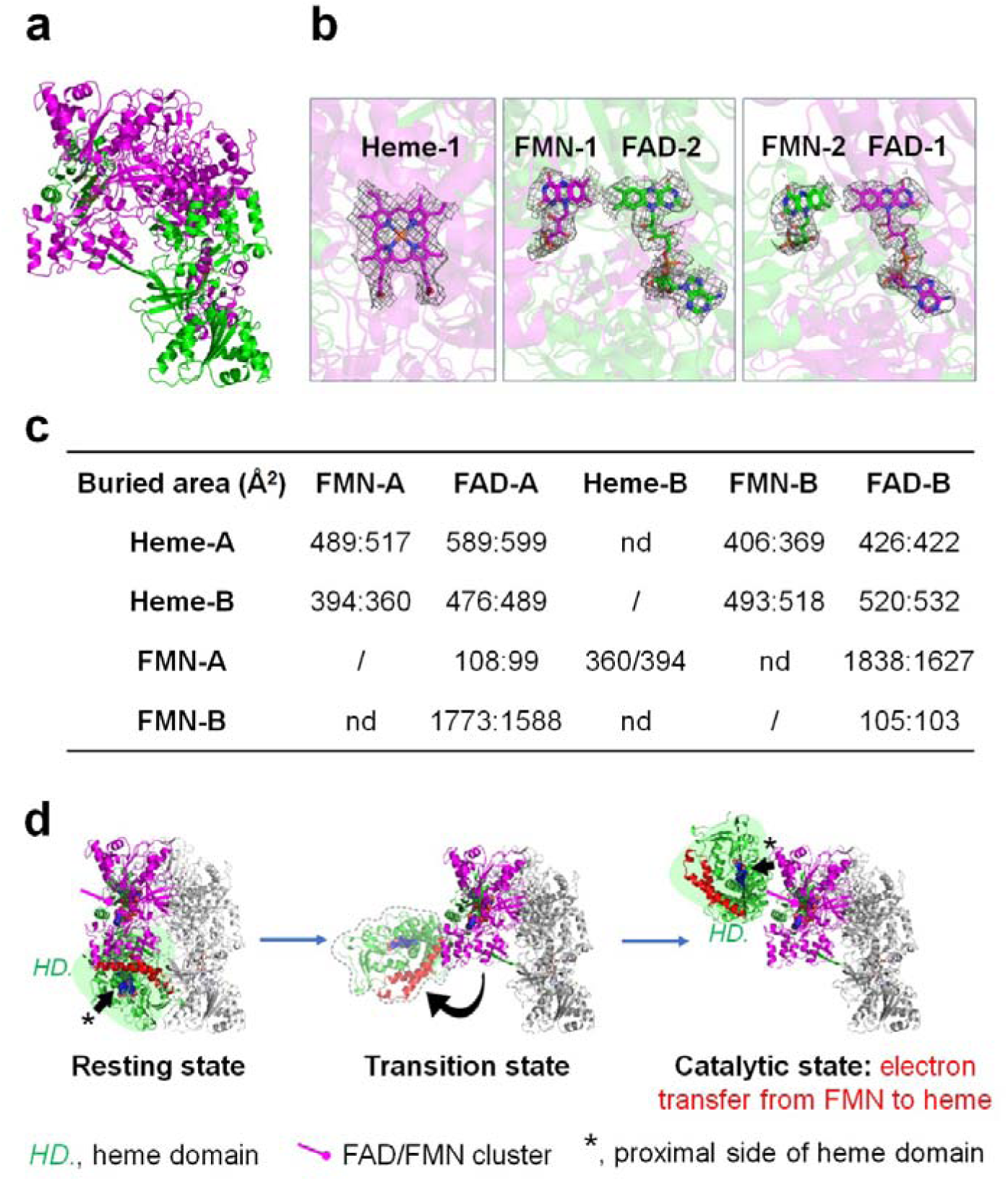
The heme-domain of *So*P450 in mobile. **a**, The cryo-EM structure of class 2 conformation of *So*P450 displayed in a cartoon model with two chains colored in green and magenta. **b**, The cofactors bound in the structure of the class 2 conformation with the cognate electron density maps displayed as described in Fig. 1b. **c**, The areas of the contact interface between each domain in the dimeric form *So*P450. nd., not detectable. **d**, The proposed electron transfer mechanism of a heme-moving model of *So*P450.

The presence of class 2 conformation implies that the heme-domain is mobile, which could be, at least partially, supported by the fact that the heme-domain forms fewer interactions to the adjacent domains than the FMN- and FAD-domain that form a CPR-domain. The buried interface of FMN-A and FAD-B is estimated to 1,838 and 1,627 Å^2^, respectively, accounting for approximately 18.5% and 7.2% of the solvent accessible surface area of each domain. The interactions between FMN-A and FAD-B, so as FMN-B and FAD-A, represent the most extensive contact among all in the dimeric configuration (**Fig. 4c**). In comparison, the heme-domain forms fewer contact to others. Accordingly, we proposed that the heme-domain is mobile and could relocate to reach the FMN-domain to acquire the electrons (**Fig. 4d**).

### The correlation of CR^I^ length and the activity of *So*P450

Based on the heme-domain moving mechanism, we built a model to simulate the catalytic state displayed in **Fig. 4d**. As shown in **Fig. 5a**, the heme-domain should flip up to allow the proximal side to approach the CPR-domain with the CR^I^-anchoring residues on either end spatially close to each other. The bulge between helix H and I of the heme-domain docks into the concave side of the CPR-domain (**Supplementary Fig. 7a**) and the contact interfaces on the heme- and CPR-domain are electrostatic complementary (**Supplementary Fig. 7b**). The edge-to-edge distance between heme-1 and FMN-1 is measured to 17.8 Å (**Fig. 5b**). The methyl moieties of FMN remains directing toward the FAD instead of being exposed to orient to the heme-domain. As previous studies reported, W574 that stacks the isoalloxazine ring of FMN assists the electron flow between FMN and heme in P450BM3^40,58^. Accordingly, the equivalent residue W591 in *So*P450 that packs against the isoalloxazine ring of FMN may serve to relay the electron from FMN (**Fig. 5b**).

**Fig. 5.**
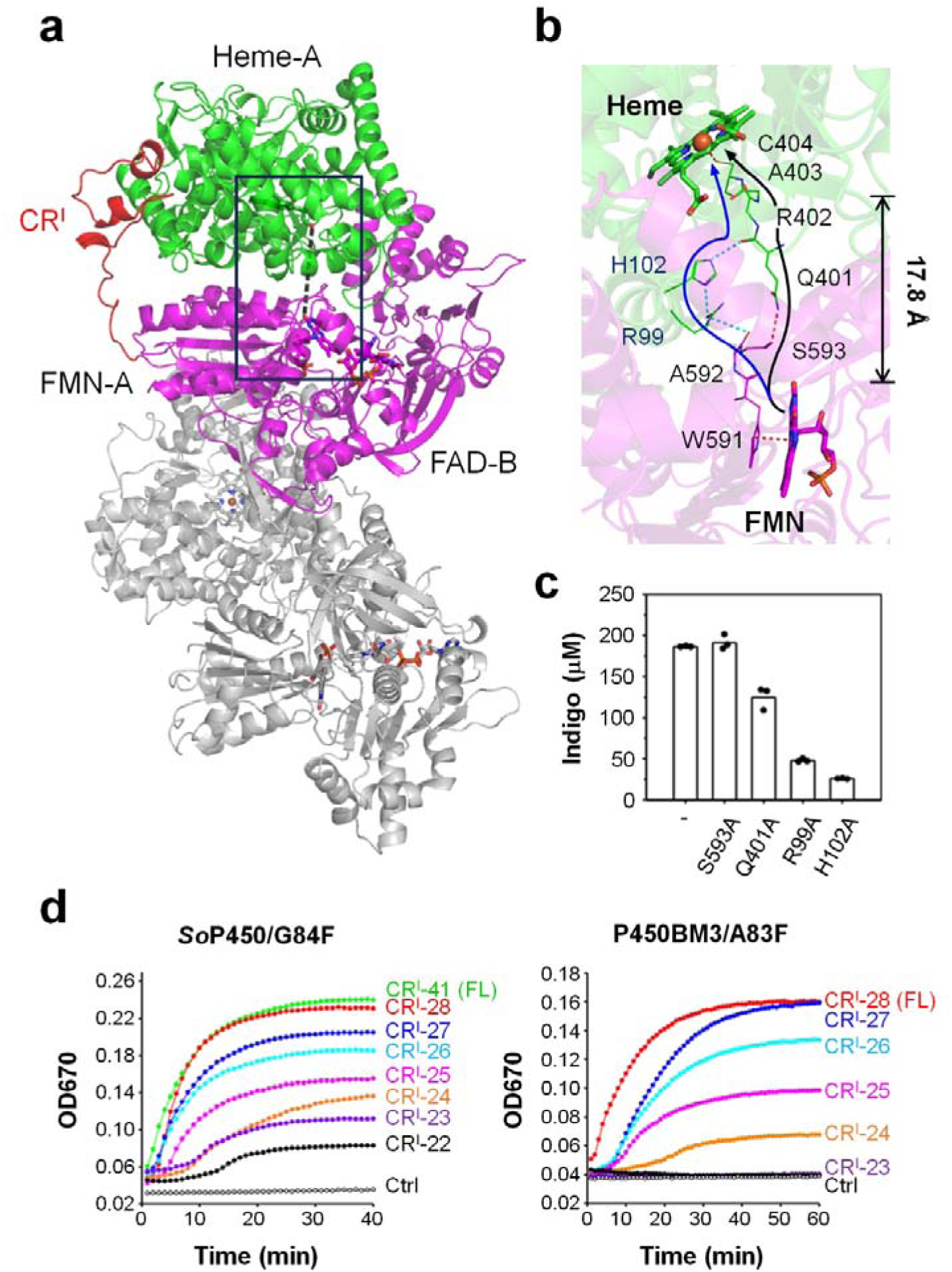
The putative electron transfer route of *So*P450. a, The model of a catalytic state of *So*P450, in which the mobile heme-domain (green) is relocated to approach the CPR-domain (magenta). The contact interface of heme- and FMN-domain is framed. The model construction processes are described in Methods. **b,** The zoom-in view of the framed area shown in **a**, with the edge-to-edge distance between heme and FMN prosthetics indicated on the right. Two possible electron traveling paths from FMN to heme are depicted by curved arrows, with nonbond jumps on the path connected by dashed lines. **c,** 1 mL reactions containing 5 µM enzyme (*So*P450/G84F and variants), 4 mM indole, and 0.2 mM NADPH were incubated at 37 °C for 20 min. The indigo production was quantified based on a standard curve established with indigo of known concentration. The average and individual values of each group are shown by bars and circles. **d,** 1 mL reactions containing *So*P450/G84F or P450BM3/A83F that possess varying length of CR^I^ assembled as described in **c** were incubated at 37 °C. The indigo formation during indicated time period was monitored via optical density at 670 nm (OD670) and the average values of a triplicate assay are displayed. The sequences of CR^I^-truncated variants are shown in **Supplementary Fig. 10**. FL, full-length; Ctrl, no enzyme. The results demonstrated in **c** and **d** are representatives of three independent experiments.

Since no prosthetic exists between FMN and heme, we suspected that the electrons shall be tunneling to the heme center through amino acids *en route* and hired HARLEM program^59^ to identify candidate residues (**Supplementary Fig. 8**). The predicted path that begins with the indole of W591 passes through covalent bonds that connect residues A592, S593, Q401, R402, A403, C404 along with a non-bonded jump between S593 O and Q401 NE2 and eventually to the heme iron (**Fig. 5b**). Mutagenesis experiment was then applied to investigate the role of these residues play in the activity of *So*P450. Because the natural substrate of *So*P450 remains unrevealed, variant G84F was constructed, based on the analogy to the variant A83F of P450BM3^60^, to confer the protein capability of synthesizing indigo by using indole as a substrate (**Supplementary Fig. 9**). Residue A592, S593, R402 and A403 contribute the main chain to the path, and C404 and W591 are indispensable to the binding of cofactors. Variant Q401A exhibits 67.1 ± 7.5 % activity of the parental enzyme, suggesting that the side chain of Q401 may play a role, but not strictly essential, in *So*P450-catalyzed reaction (**Fig. 5c**). This suggests that the electrons may bifurcate to another path such as one that comprises W591, A592, R99, H102 and then to the main chain of Q401 (**Fig. 5b**). Indeed, substituting R99 and H102 with Ala reduced the enzyme activity to 48 ± 1.9 % and 24.6 ± 0.4 %, respectively (**Fig. 5c**).

This model also suggests that the CR^I^ fragment of adequate length shall be required for the heme-domain to reach for the CPR-domain. Therefore, we constructed a series of variants with the CR^I^ progressively truncated and measured their activity. Variant *So*P450/G84F-CR^I^-28 that retains 28 amino acids in the CR^I^ segment, the same as that of P450BM3 (**Supplementary Fig. 10**), exhibits activity comparable to the full-length protein (**Fig. 5d**). Further truncation led to progressive reduction in enzyme activity, suggesting that shorter CR^I^ might constrain the optimal docking of heme-domain to the CPR-domain and that our model of a catalytic state of *So*P450 may be reliable. This is also the case for P450BM3, that the catalytic activity decreased as the CR^I^ was shortened (**Fig. 5d**). Intriguingly, the length of CR^I^ fragment of P450s tested in our study ranges from 26 (*Mt*P450 from *Marinactinospora thermotolerans*) to 41 amino acids (*So*P450) (**Extended Data Table 1**), and the shortest CR^I^ that can be identified through a searching in the GenBank consists of 25 amino acids (**Supplementary Fig. 11**). Altogether, 25 amino acids could be the minimal requirement for the CR^I^ in P450BM3 homologous CPR-containing self-sufficient P450s to exert the optimal activity.

## Conclusion

In this study, the cryo-EM and crystal structures of *So*P450 were resolved to reveal the unique dimeric organization of a P450BM3 homologous CPR-containing self-sufficient P450. The identification of the class 2 conformation that lacks one heme-domain suggests that the heme-domain is mobile and might relocate to reach the CPR-domain for electron acquisition. Accordingly, the catalytic state was proposed through modeling study, which reveals putative electron transfer paths from FMN to heme. Finally, the extensive CR^I^ segment that connects the heme- and FMN-domain should govern the transition of the heme-domain between the resting and the catalytic state. While the association of the length of CR^I^ segment and the catalytic activity is demonstrated, the correlation of other properties of this region (e.g., rigidity) remains to be explored. Given the high protein sequence identity between *So*P450 and P450BM3, the reported structures would be a reliable model for predicting the three-dimensional structure of the latter. Intriguingly, the indigo synthesis assay based on the Phe variant indicates that *So*P450 exhibits higher catalytic activity than P450BM3, illuminating further application potentials of *So*P450. These results are of fundamental importance to advance our understanding about the molecular basis of P450BM3 as well as other homologous P450s, but provide valuable information for engineering and applications of these enzymes.

## Data availability

All data generated or analyzed during this study are included in the manuscript and the Supplementary Information files. The 3D cryo-EM density maps of *So*P450 (EMD-66257), *So*P450-class 2 (EMD-66258), *So*P450/NADPH (EMD-66267) and *So*P450-CR^I^-10 (EMD-66262) have been deposited in the Electron Microscopy Data Bank (EMDB) database. The atomic model by X-ray crystallography *So*P450 (PDB ID, 9X0N) has been deposited in the Protein Data Bank (PDB).

## Methods

### Plasmid construction and mutagenesis

The genes that encode CPR-containing self-sufficient P450s containing His-tag, tobacco etch virus (TEV) protease cleavage site (ENLYFQG) and (AG)*_n_* linker on the *N*-terminus were chemically synthesized and cloned into expression vector pET-46 Ek/LIC. The amino acid sequences of all selected P450s are listed in **Supplementary Table 1**. The polymerase chain reaction-based site-directed mutagenesis was conducted to construct variants by using plasmid that encodes *So*P450 or P450BM3 as a template with mutagenic oligonucleotides listed in **Supplementary Table 2**. All plasmids were verified by direct sequencing.

### Protein expression and purification

For protein expression, the plasmid was transformed into *Escherichia coli* BL21 (DE3). The transformants were initially cultured overnight in 100 mL Luria–Bertani (LB) medium at 37 °C containing 100 µg mL^-1^ ampicillin. The preculture was inoculated into 5 L LB medium containing 100 µg mL^-1^ ampicillin and grown at 37 °C with constant shaking at 220□rpm. Protein expression was induced when the optical density at 600 nm (OD600) reached 0.6 by adding 0.3 mM IPTG (isopropyl β-D-1-thiogalactopyranoside) and 5 µM chlorhematin, followed by cultivation at 16 °C for 18 h. The cells were collected by centrifuging at 6,000 rpm for 10 min, and resuspended in 100□mL of lysis buffer containing 25□mM Tris-HCl (pH□7.5), 150□mM NaCl, and 20□mM imidazole. Cell disruption was achieved using a French press (JuNeng Biology & Technology), followed by centrifugation at 17,000 rpm for 60 min to pellet the debris. The supernatant was then loaded onto a Ni-NTA column using a protein chromatography system (Sepure Instruments, Suzhou, China) and eluted with a gradient of imidazole ranging from 20 to 500 mM. The fractions containing target protein were collected and dialyzed against a 25 mM HEPES buffer (pH 7.5) containing 150 mM NaCl. The proteins were supplemented with 10 mM DTT and then passed through a Superdex 200 10/300 GL column (GE Healthcare, Madison, WI). The peak fractions were collected, quantified and concentrated for further analyses.

### Cryo-EM sample preparation and data collection

Prior to the sample preparation, the Electron Microscopy Sciences grids (CFlat, Au, R 1.2/1.3) were discharged with 15 mA at 0.39 mbar for 45 s. 4 μL of protein solution (1 mg mL^-1^) was applied to the grids, and 10 mM NADPH was supplemented for the preparation of the complex of *So*P450/NADPH. A Vitrobot Mark IV (Thermo Fisher Scientific) was used to blot the grid for 3 s at 4 °C and 100% humidity, followed by plunge-freezing into liquid ethane cooled by liquid nitrogen. The grids were then transferred into a box stored in liquid nitrogen before image acquisition. The datasets were collected with a Titan Krios electron microscope equipped with a Falcon 4 detector by using the EPU software (Thermo Fisher Scientific). The data collection parameters are outlined in **Extended Data Table 2**.

### Cryo-EM data processing

All datasets were processed using cryoSPARC v.4.6.0^64^, with the details outlined in **Extended Data Fig. 1, 4 and 5**. In general, all movie frames underwent motion correction and contrast transfer function (CTF) prior to an automated particle picking. After multiple rounds of picking, ultimately yielded particles were subjected to ab-initio reconstruction with C1 symmetry imposed. After heterogeneous refinement to optimized data classification, select classes with high quantity for homogeneous refinement and non-uniform refinement. Further refined the correction of particle motion trajectories by reference-based motion correction, and repeated homogeneous refinement and non-uniform refinement were performed. For *So*P450, the final reconstructions were obtained at resolutions of 2.72 Å from a set of 740,228 particles (class 1 conformation, the dimer form) and 3.08 Å from a set of 154,436 particles (class 2 conformation, the one-heme-missing form) after local refinement. The resolution of the final maps was estimated using the gold-standard Fourier shell correlation (FSC) with a 0.143 criterion.

For *So*P450-CR^I^-10, 4,826 movie stacks were collected and similarly processed. 4,684,184 particles were auto-picked and extracted from the preprocessed micrographs. 1,407,631 particles were selected for 3D reconstruction, and 1,180,657 particles were kept for producing the final density map determined at a resolution of 2.34 Å.

For *So*P450/NADPH, 2,754 movies were acquired. The movie stacks were motion corrected using patch motion correction with a binning factor of 2 for further data processing (final pixel size of 0.926□Å on the sample level), and CTF estimation was performed with patch CTF estimation. After auto-picking, 437,136 particles were extracted for 2D classification, from which 139,104 particles were retained for reconstruction, yielding a density map at 2.68 Å resolution.

### Cryo-EM structure determination

The initial models of each domain of *So*P450 were obtained by using AlphaFold3 algorithm^65^. These models were docked into the density map using UCSF Chimera^66^. The structure models were then refined against the sharpened maps with Phenix real-space refine^67^, ISOLDE^68^ and Coot^69^. The models used for *So*P450-CR^I^-10, *So*P450-NADPH complex and the class-2 conformation of *So*P450 were constructed based on the dimeric *So*P450 model. Summary statistics for map reconstruction and model building are presented in **Extended Data Table 2**.

### Crystallization, structure determination and refinement

The initial crystallization screening for SoP450 was performed at 22 □ by the sitting-drop vapor-diffusion. Protein solution (1 µL) was mixed with 1 µL of reservoir solution and equilibrated against 100 µL of reservoir solution. Needle-shaped reddish-brown crystals were observed under the condition containing 0.1 M HEPES (pH 7.0), 15% PEG 20,000. After optimizing the crystallization conditions, reddish-brown strip-shaped crystals were obtained under 0.1 M HEPES (pH 7.5), 15% PEG 20,000. Prior to X-ray diffraction data collection, crystals were cryoprotected by soaking in a buffer containing 0.1 M HEPES (pH 7.5) and 20% PEG 20,000. The X-ray diffraction datasets were collected at beam line TPS 05A of the National Synchrotron Radiation Research Center (NSRRC, Hsinchu, Taiwan). Data was processed by using HKL2000^70^.

The crystal structure of *So*P450 was solved by molecular replacement using the Phaser program^71^ by using the cryo-EM structure of the dimeric *So*P450 as a search model. Prior to structure refinement, 5% of the reflections were randomly selected and set aside for calculating *R*_free_ as a monitor of model quality. The model was manually adjusted using Coot^69^ and refined with Refmac5 in CCP4 suit^72^ and PHENIX^67^. The statistics of data collection and refinement are summarized in **Extended Data Table 3**.

### Enzyme activity measurement

For indigo formation assay, the reaction mixture (1 mL) containing purified enzyme (5 μM) and indicated concentration of indole were added in 0.1 M phosphate buffer (pH 8.0). The reaction was initiated by adding NADPH to a final concentration of 0.2 mM. The formation of indigo at 37 □ was monitored by measuring the absorbance at 670 nm at destined timepoints.

### Model preparation

The reconstruction of CR^I^ segment missing in the resolved structures of *So*P450 was performed by using the loop modeling module implemented in Rosetta 3.15^73^. In the absence of available homologous templates, the CR^I^ segment was built de novo using the cyclic coordinate descent algorithm, which relies purely on physical principles^74^. To account for conformational diversity, at least 128 distinct conformations were generated for each loop region requiring completion.

To model the relocated heme-domain approaching the FMN-domain (a “catalytic state”), protein–protein docking was performed using the local docking protocol in Rosetta 3.15. The most plausible pose—in which the heme-domain was positioned close to the FMN-domain to enable efficient electron transfer, and the loop-anchoring residues were spatially proximal to the heme-domain anchor points, thereby ensuring appropriate loop accessibility—was manually selected for subsequent loop modeling and molecular dynamics simulations.

### Molecular dynamics simulations

The CR^I^-containing models and the relocated heme–FMN complex were subjected to molecular dynamics (MD) simulations using the AMBER22 software package^75^, with three independent replicas performed for each system to ensure reproducibility. During the MD process, the Amber FF14SB^76^ force field was used for the protein and the TIP3P^77^ model was employed for the solvent waters. The heme cofactor parameters were generated using the Metal Center Parameter Builder (MCPB) approach^78^. The restrained electrostatic potential (RESP)^79^ charges of all ligands were calculated at the HF/6-31G* level using the Gaussian 16 package^80^, and the ligands were subsequently described using the AMBER GAFF□ force field, consistent with previous MD studies on cytochrome P450 systems^81–84^. All simulations were conducted under periodic boundary conditions, with cubic solvent boxes applied to each system. The initial coordinates and topology files were generated using the tleap module implemented in AMBER22. Sodium or chloride ions were added as needed to neutralize the total system charge. In total, each system comprised approximately 220,000 atoms. Energy minimization was performed in three stages: first on the solvent molecules, then on the protein side chains, and finally on all atoms. Each minimization stage consisted of 4000 cycles of steepest descent followed by 2000 cycles of conjugate gradient minimization. After minimization, each system was gradually heated from 0 K to 300 K under the NVT ensemble (with Langevin thermostat)^85^, followed by a 100 ps NPT ensemble density equilibration at 300 K and 1.0 atm (with Berendsen barostat)^86^. Subsequently, 100 ns production runs were carried out for each system—under the NVT ensemble with a 1 fs time step. A nonbonded cutoff of 10 Å was applied to both van der Waals (Lennard–Jones 12-6) and real-space electrostatic interactions, with long-range electrostatics treated by the particle mesh Ewald (PME) method^87^. The SHAKE^88^ algorithm was applied to constrain all bonds involving hydrogen atoms, thereby removing the high-frequency stretching vibrations of X–H bonds during the MD simulations. To investigate the electron transfer chain, conformational clustering was performed for the relocated heme–CPR complex model based on protein backbone coordinates. The most populated cluster accounted for 83.7% of the total conformational ensemble (**Fig. 5a** and **5b**). Trajectory analyses, including clustering, RMSD, and RMSF calculations, were carried out using the parallelized cpptraj module implemented in the AMBER 22 suite^89,90^.

## Supporting information

Supplementary Tables 1 to 2 Supplementary Figs. S1 to S13 Supplementary Computation Information References

## Acknowledgments

This work was supported by the National Key Research and Development Program of China (2021YFC2104000), Hubei Hongshan Laboratory (2022hszd030), the National Natural Science Foundation of China (32371307, 32271318, 82341210, 22473118 and 82430108); and the Interdisciplinary Research Project of Hangzhou Normal University (2024JCXK02). We thank NSRRC (National Synchrotron Radiation Research Center, Taiwan) for access to beam lines TPS-05A and TPS-07A that contributed for the synchrotron data collection.

---

**Extended Data Fig. 1.**
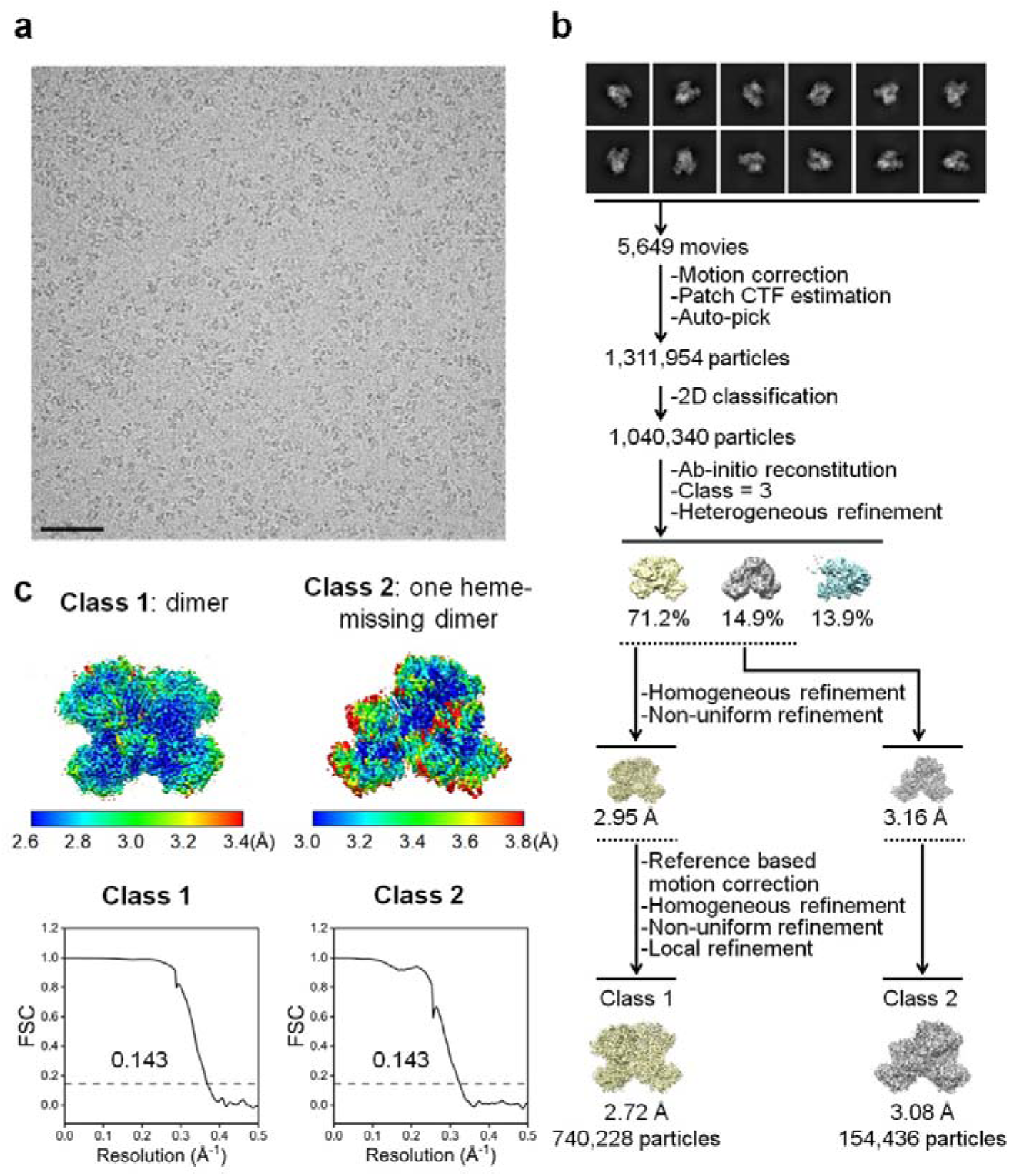
Workflow for image processing and structure reconstruction of cryo-EM analyses of *So*P450. **a**, A representative micrograph of *So*P450. Scar bar, 50 nm. **b**, Reference free 2D averages and cryo-EM maps at the various stages of processing. **c**, Local resolution estimation of the finalized cryo-EM maps and corrected curve of the global Fourier shell correlation (FSC) with 0.143 gold-standard criterion.

**Extended Data Fig. 2.**
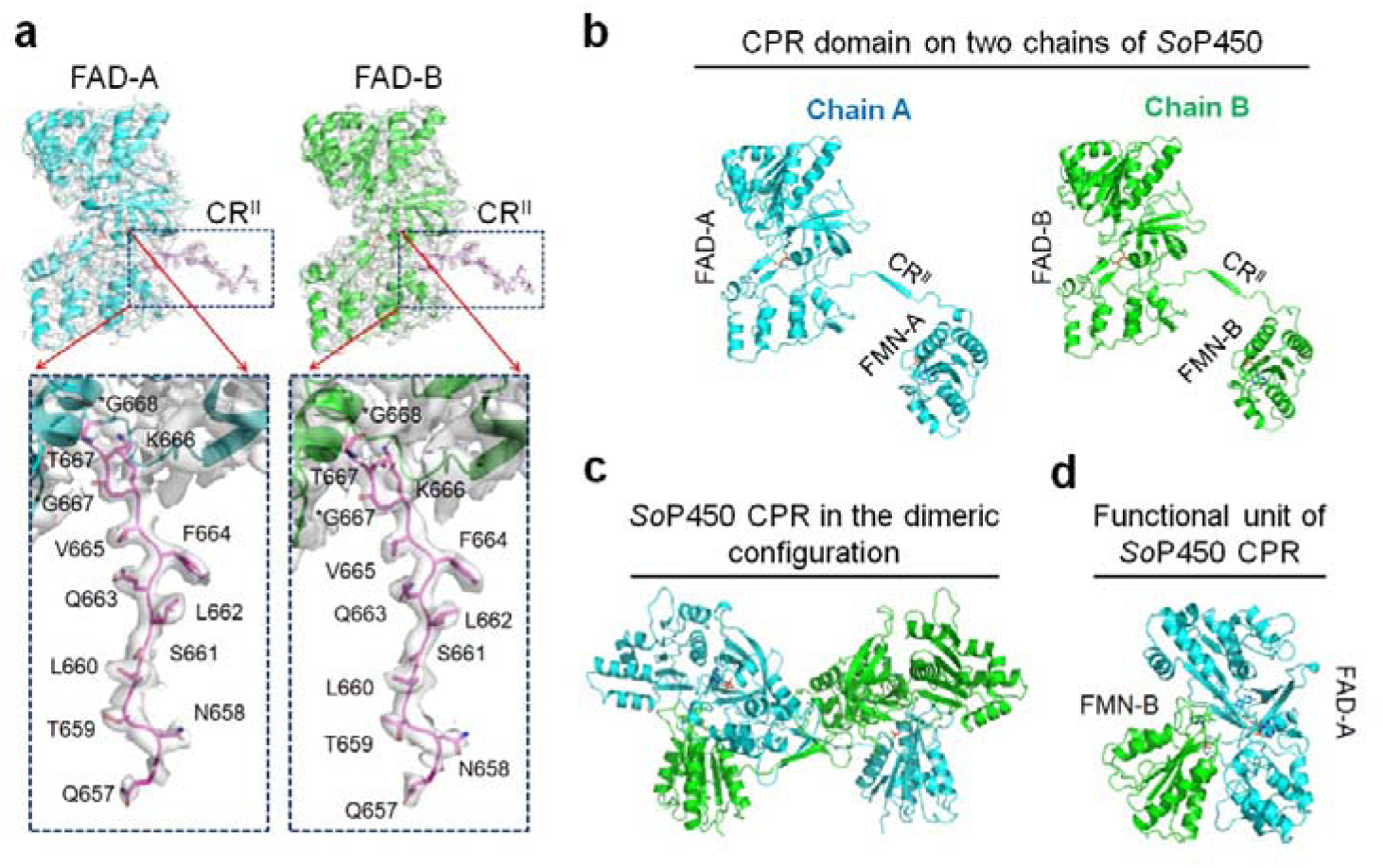
The assignment of flavin-containing domains of *So*P450. **a**, The cartoon models, amino acids and electron density maps of CR^II^ contoured at 5.0 σ of the cryo-EM structure of *So*P450 (PDB ID, 9WUC). **b**, The cartoon models of FMN- and FAD-domains of *So*P450 in two individual polypeptide chains. **c**, The CPR of *So*P450 in the dimeric configuration. **d**, One functional CPR-domain of *So*P450.

**Extended Data Fig. 3.**
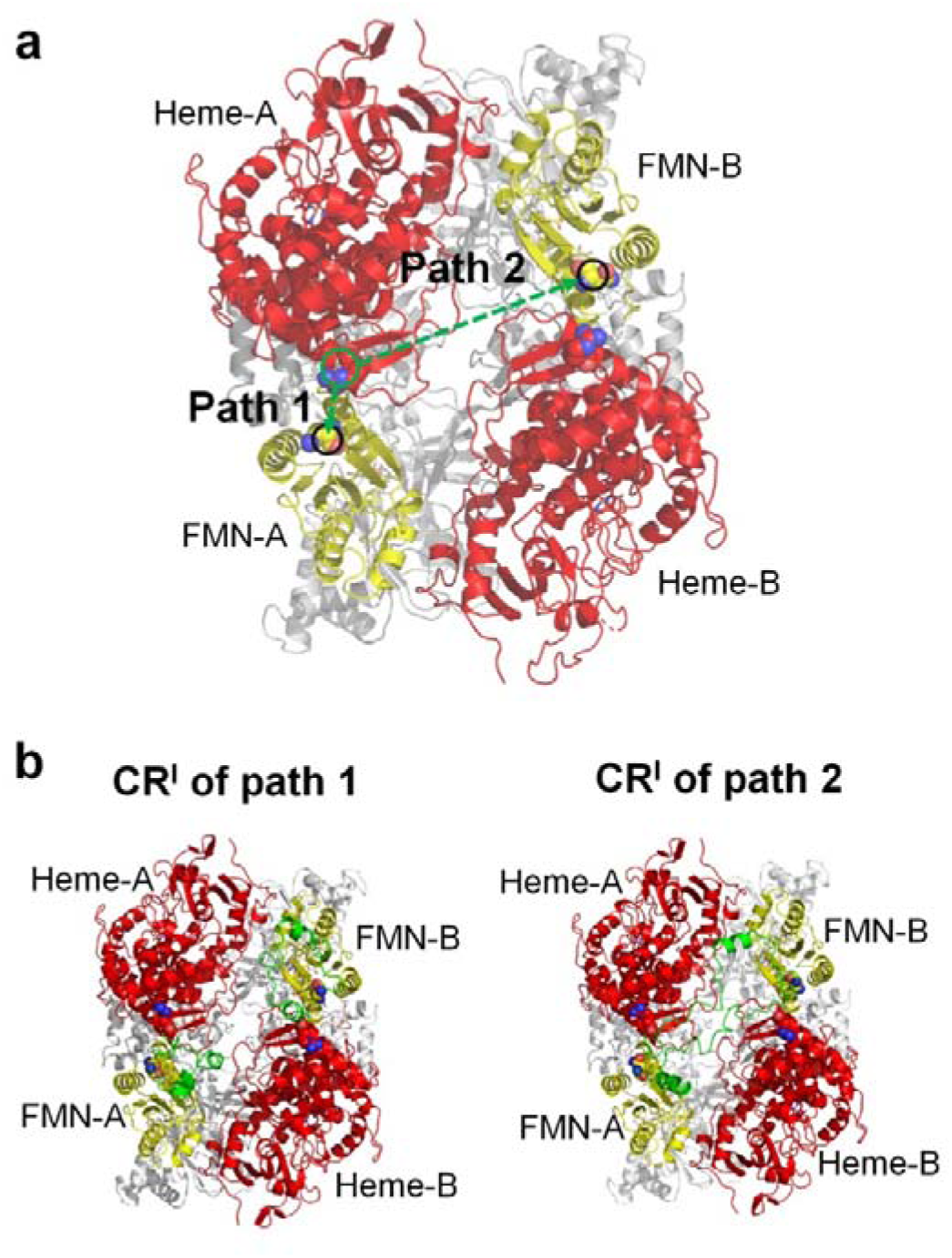
Two possible models based on the direction of the CR^I^ segment. **a**, The cartoon model of cryo-EM structure of *So*P450 (PDB ID, 9WUC) with two paths that connect heme-A to FMN-A or FMN-B indicated with green dashed lines. Green circle, the C-terminus of heme-A; black circles, the N-termini of FMN-A and FMN-B. **b**, The models based on two possible paths, with CR^I^ fragments colored in green. The model construction processes are described in Supplementary Computation Information and **Supplementary Fig. 12**.

**Extended Data Fig. 4.**
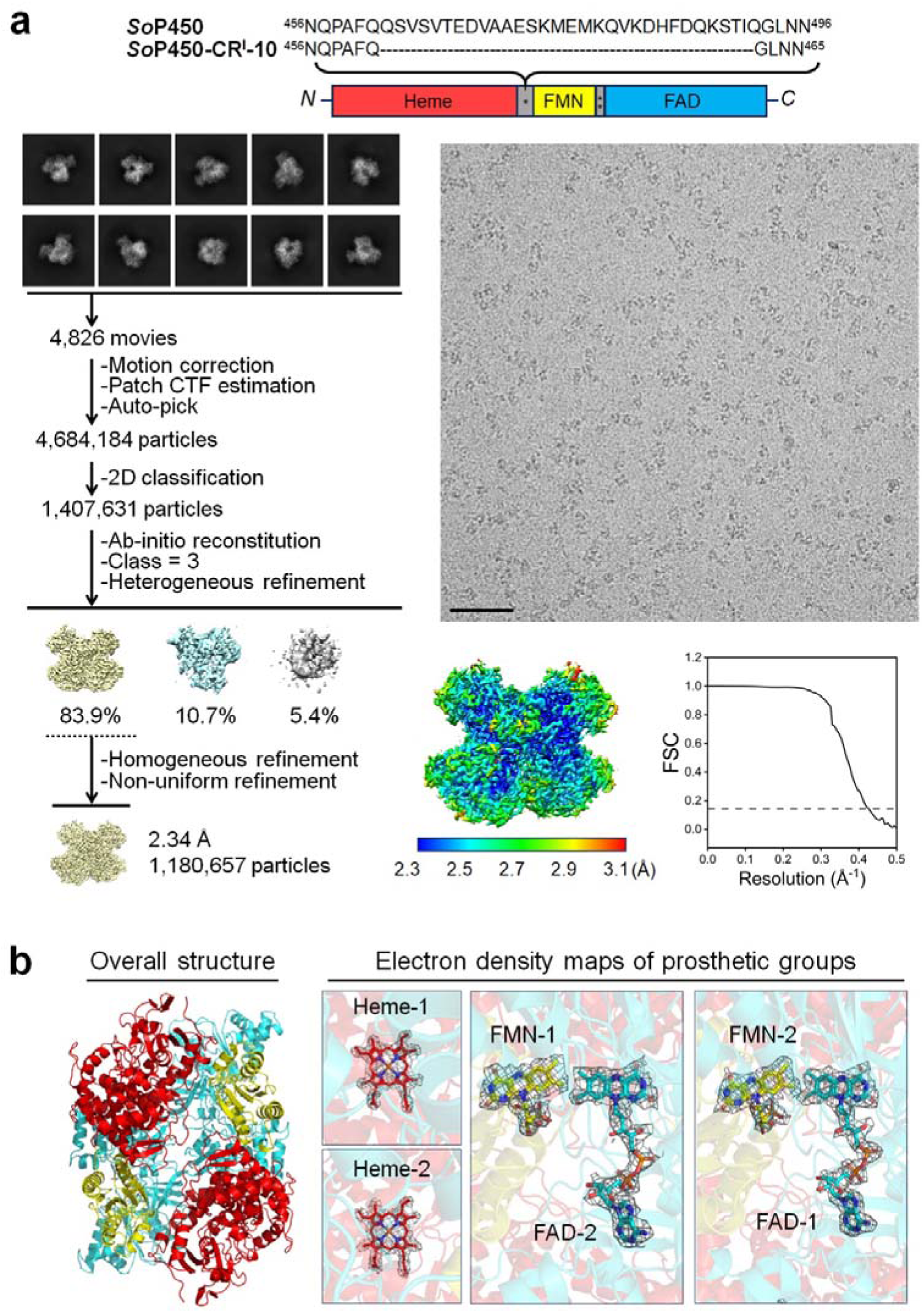
Workflow for image processing and structure reconstruction of cryo-EM analyses of *So*P450-CR^I^-10. **a**, A representative micrograph, reference free 2D averages, cryo-EM maps at the various stages of processing, local resolution estimation of the finalized cryo-EM maps and corrected curve of the global FSC with 0.143 gold-standard criterion. Scar bar, 50 nm. Top, the amino acid sequences of the full-length (*So*P450) and CR^I^-truncated variant (*So*P450-CR^I^-10) of *So*P450. **b**, The overall structure of *So*P450-CR^I^-10 (PDB ID, 9WUK) and electron density maps of prosthetic groups bound in the structure contoured at 5.0 σ are shown as described in Fig. 1b.

**Extended Data Fig. 5.**
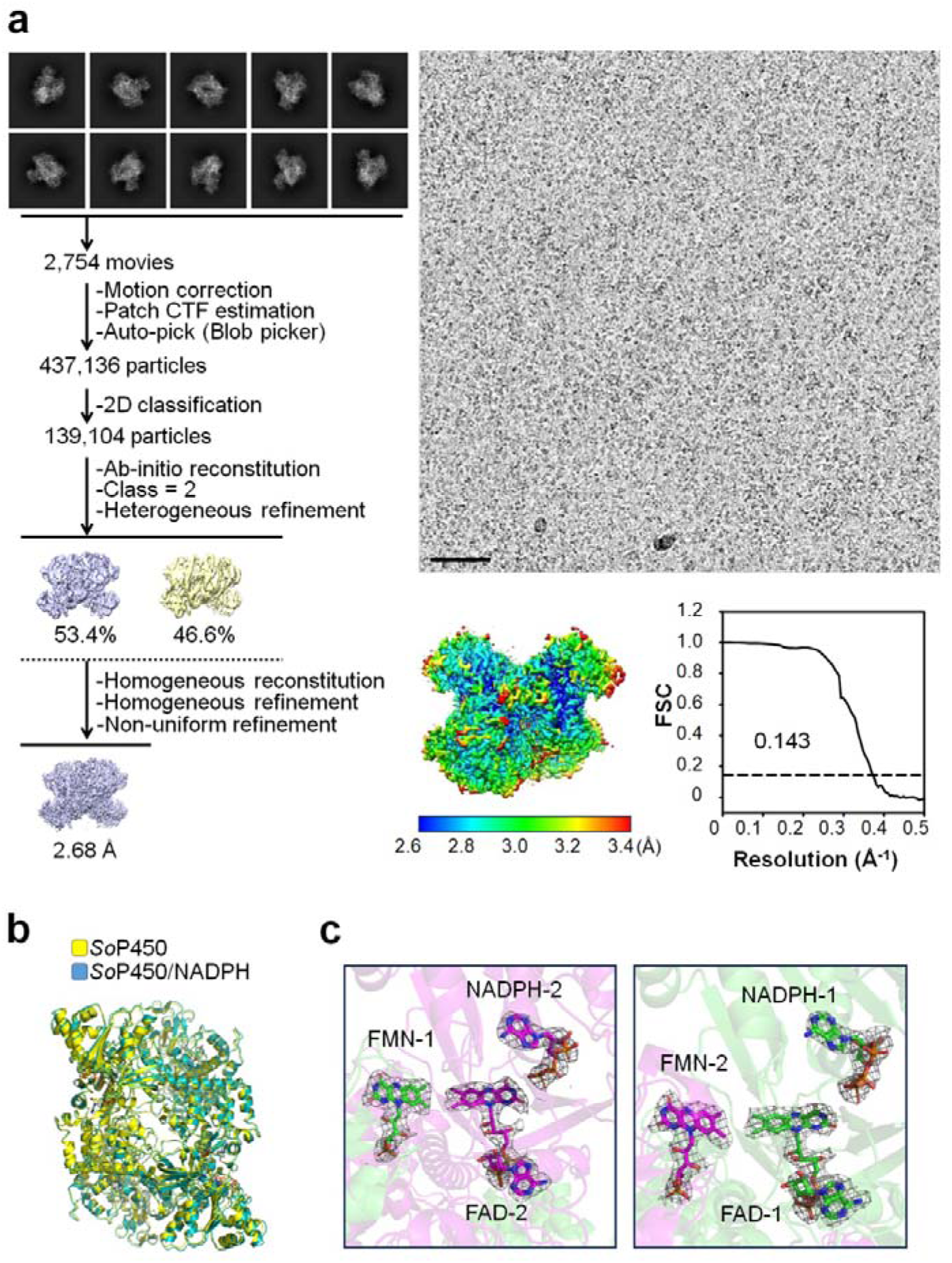
Workflow for image processing and structure reconstruction of cryo-EM analyses of of *So*P450/NADPH. **a**, A representative micrograph, reference free 2D averages, cryo-EM maps at the various stages of processing, local resolution estimation of the finalized cryo-EM maps and corrected curve of the global FSC with 0.143 gold-standard criterion of the complex of *So*P450/NADPH (PDB ID, 9WUP). Scar bar, 50 nm. **b**, The structure superimposition of *So*P450 and *So*P450/NADPH. **c**, The electron density maps of prosthetic groups bound in *So*P450/NADPH contoured at 5.0 σ are shown as described in Fig. 1b.

**Extended Data Fig. 6.**
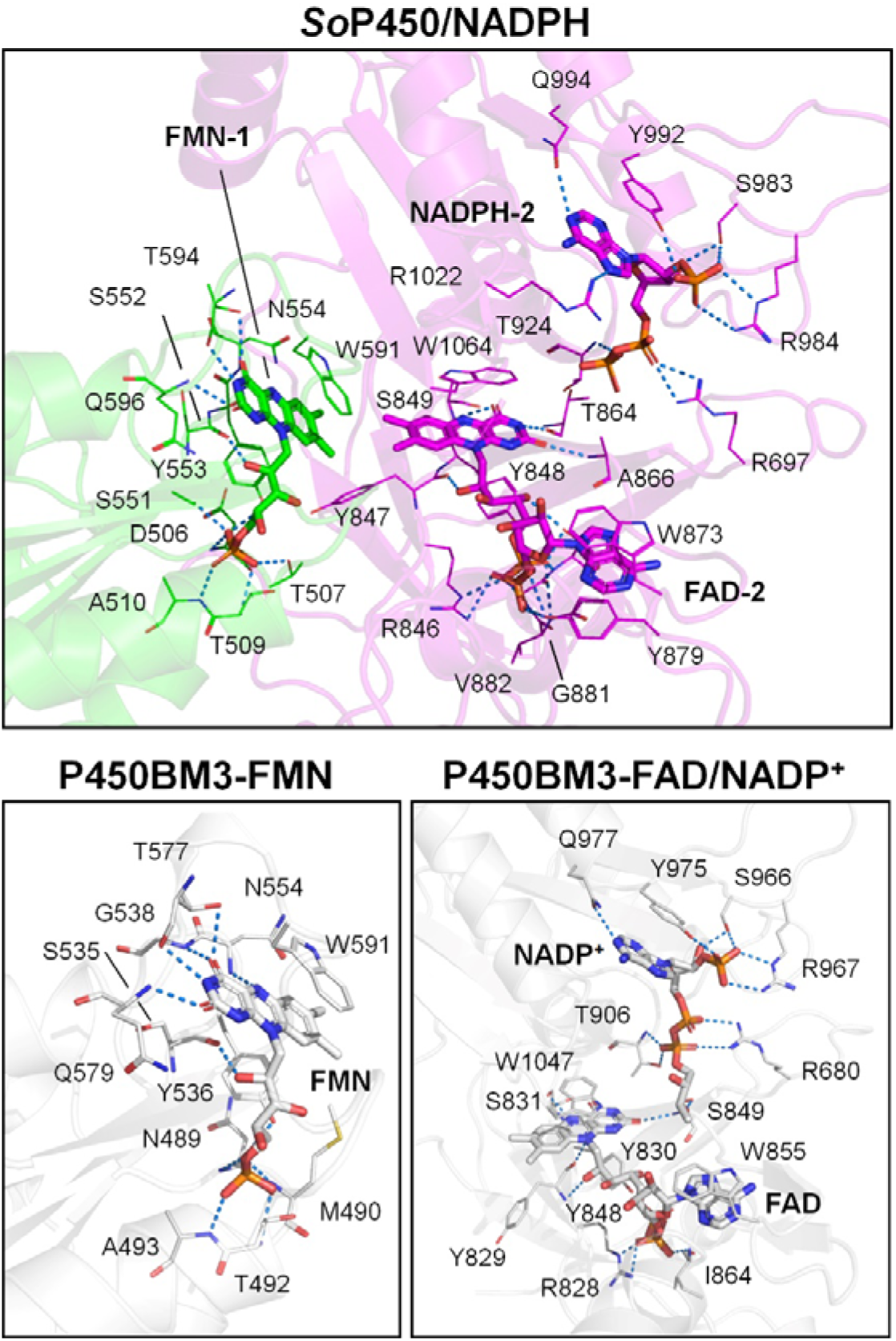
Interaction network of FMN, FAD and NADPH in *So*P450/NADPH (PDB ID, 9WUP) and P450BM3. Protein residues and cofactors are displayed in line and sticks, respectively. Proteins belong to two individual chains in the homodimer are colored in green and magenta. The drawings of P450BM3-FMN and P450BM3-FAD/NADP^+^ are based on PDB entry 1BVY and 4DQL, respectively. Dashed lines, distance < 3.5 Å.

**Extended Data Table 1.** Basic information of CPR-containing self-sufficient P450s tested in this study.

| Organism | GenBank no. | Identity to P450BM3 | Total residue | CR<sup>I</sup> length* | Ref. |
| --- | --- | --- | --- | --- | --- |
| *Bacillus megaterium* | WP_034650526 | 100% | 1048 | 28 | 30 |
| *Shimazuella soli* | WP_240877128 | 60.4% | 1067 | 41 | This study |
| *Thermothelomyces thermophilus* ATCC 42464 | XP_003663647 | 35.5% | 1080 | 42 | 61 |
| *Marinactinospora thermotolerans* | WP_078760863.1 | 40.6% | 1039 | 26 | 62 |
| *Thermocatellispora tengchongensis* | WP_312924648.1 | 40% | 1030 | 30 |  |
| *Streptomyces cattleya* | WP_014145542.1 | 39.9% | 1068 | 31 |  |
| *Aspergillus thermomutatus* | XP_026611160 | 36.2% | 1020 | 28 |  |
| *Pseudonocardia thermophila* | WP_073455647.1 | 42.6% | 1054 | 28 |  |
| *B. nakamurai* | WP_061522868 | 58.9% | 1052 | 28 |  |
| *B. cereus* | WP_000412767 | 60.1% | 1065 | 37 | 63 |

\*The number of amino acids.

**Extended Data Table 2.** Cryo-EM data collection, refinement and validation statistics.

|  | *SoP450* | *SoP450-clas*<br>*s 2* | *SoP450/*<br>*NADPH* | *SoP450-CR<sup>I</sup>*<br>*-10* |
| --- | --- | --- | --- | --- |
| **PDB entry** | 9WUC | 9WUD | 9WUP | 9WUK |
| **EMDB-entry** | EMD-6625<br>7 | EMD-6625<br>8 | EMD-66267 | EMD-6626<br>2 |
| **Data collection and processing** |  |  |  |  |
| Magnification | 130,000 | 130,000 | 130,000 | 130,000 |
| Voltage (kV) | 300 | 300 | 300 | 300 |
| Detector | Falcon 4 | Falcon 4 | Falcon 4 | Falcon 4 |
| Electron exposure (e<sup>-</sup>/Å<sup>2</sup>) | 52 | 52 | 50 | 52 |
| Defocus range (μm) | -1.0 to -2.0 | -1.0 to -2.0 | -1.4 to -1.6 | -1.0 to -2.0 |
| Pixel size (Å) | 0.93 | 0.93 | 0.463 | 0.93 |
| Symmetry imposed | C1 | C1 | C1 | C1 |
| Initial particle images (no.) | 1,311,954 | 1,311,954 | 437,136 | 4,684,184 |
| Final particle image (no.) | 740,228 | 154,436 | 139,104 | 1,180,657 |
| FSC threshold | 0.143 | 0.143 | 0.143 | 0.143 |
| Map resolution (Å) | 2.72 | 3.08 | 2.68 | 2.34 |
| **Refinement** |  |  |  |  |
| Initial model used | de novo | *SoP450* | *SoP450* | *SoP450* |
| Model resolution (Å) | 3.02 | 3.53 | 3.03 | 2.73 |
| FSC threshold | 0.5 | 0.5 | 0.5 | 0.5 |
| Map sharpening *B* factor (Å<sup>2</sup>) | 103.3 | 88.5 | 72.1 | 76.8 |
| **Model composition** |  |  |  |  |
| Non-hydrogen atoms | 16628 | 12894 | 16672 | 16628 |
| Protein residues | 2050 | 1598 | 2048 | 2050 |
| Ligands | 6 | 5 | 8 | 6 |
| ***B* factors (Å<sup>2</sup>)** |  |  |  |  |
| Protein | 49.93 | 79.62 | 59.56 | 55.1 |
| Ligand | 22.44 | 54.87 | 41.09 | 30.74 |
| **R.m.s. deviations** |  |  |  |  |
| Bond lengths (Å) | 0.003 | 0.004 | 0.004 | 0.002 |
| Bond angles (°) | 0.580 | 0.624 | 0.932 | 0.516 |
| **Validation** |  |  |  |  |
| MolProbity score | 1.75 | 1.68 | 1.38 | 1.61 |
| Clashscore | 9.95 | 9.79 | 5.99 | 7.58 |
| Poor rotamers (%) | 0.45 | 0.74 | 0.45 | 0.62 |
| **Ramachandran plot** |  |  |  |  |
| Favored (%) | 96.47 | 97.05 | 97.79 | 96.87 |
| Allowed (%) | 3.53 | 2.95 | 2.11 | 3.13 |
| Outliers (%) | 0 | 0 | 0.1 | 0 |

**Extended Data Table 3.**
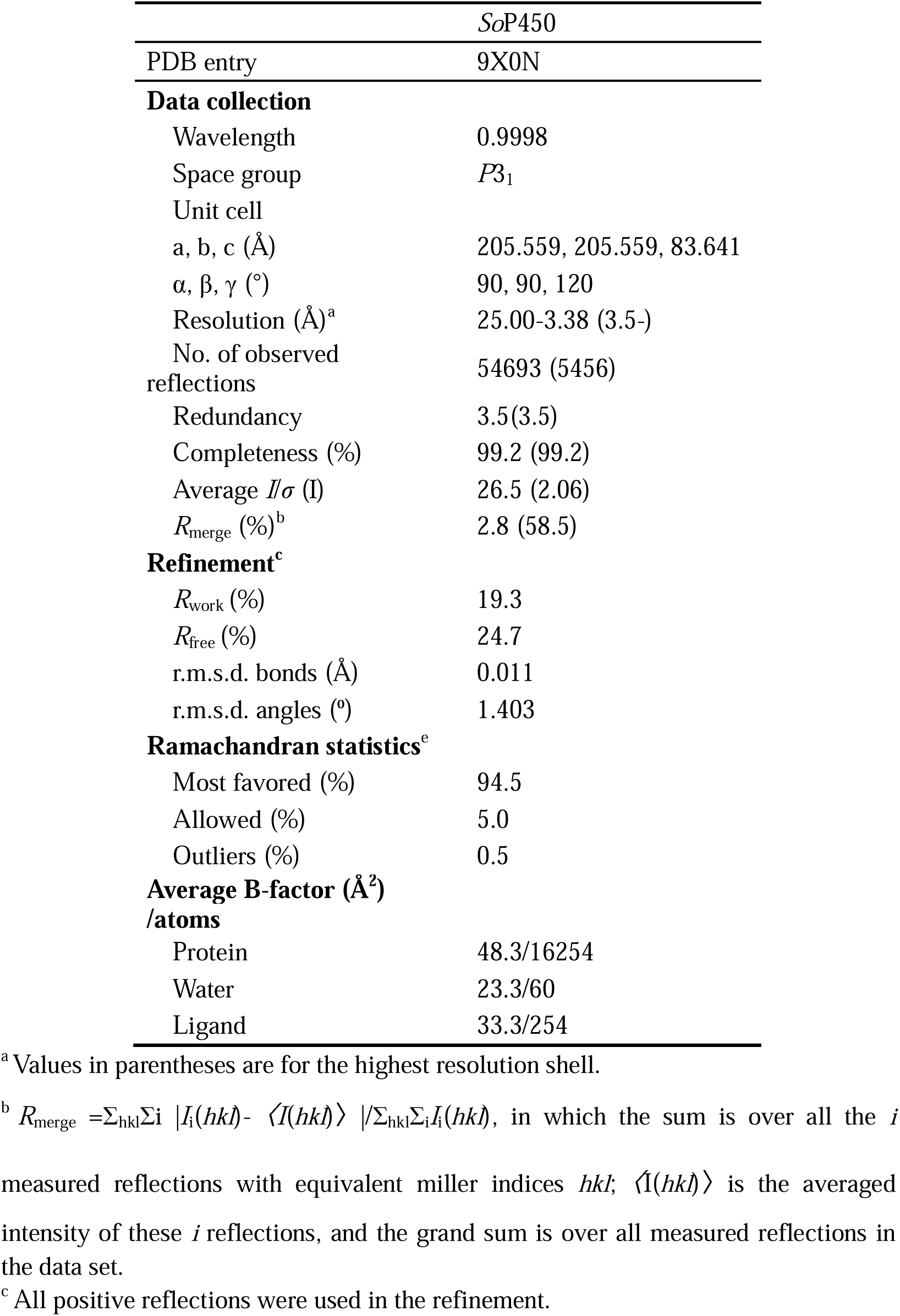
Data collection and refinement statistics of crystal structure of *So*P450.

