## Supplementary Tables 1 to 2 Supplementary Figs. S1 to S13 Supplementary Computation Information References for "Full-length structure of CPR-containing self-sufficient cytochrome P450"

**This PDF file includes**

Supplementary Tables 1 to 2

Supplementary Figs. S1 to S13

Supplementary Computation Information

References

**Supplementary Table 1. Amino acid sequences of P450s in Extended Data Table 1**

*Bacillus megaterium* P450 (P450BM3)

HHHHHHHHHHHHENLYFQGAGSGAGAGAGIKEMPQPKTFGELKNLPLLNTDKPVQALMKIADELGEIFKFEAPGRVTRYLSSQRLIKEACDESRFDKNLSQALKFVRDFAGDGLFTSWTHEKNWKKAHNILLPSFSQQAMKGYHAMMVDIAVQLVQKWERLNADEHIEVPEDMTRLTLDTIGLCGFNYRFNSFYRDQPHPFITSMVRALDEAMNKLQRANPDDPAYDENKRQFQEDIKVMNDLVDKIIADRKASGEQSDDLLTHMLNGKDPETGEPLDDENIRYQIITFLIAGHETTSGLLSFALYFLVKNPHVLQKAAEEAARVLVDPVPSYKQVKQLKYVGMVLNEALRLWPTAPAFSLYAKEDTVLGGEYPLEKGDELMVLIPQLHRDKTIWGDDVEEFRPERFENPSAIPQHAFKPFGNGQRACIGQQFALHEATLVLGMMLKHFDFEDHTNYELDIKETLTLKPEGFVVKAKSKKIPLGGIPSPSTEQSAKKVRKKAENAHNTPLLVLYGSNMGTAEGTARDLADIAMSKGFAPQVATLDSHAGNLPREGAVLIVTASYNGHPPDNAKQFVDWLDQASADEVKGVRYSVFGCGDKNWATTYQKVPAFIDETLAAKGAENIADRGEADASDDFEGTYEEWREHMWSDVAAYFNLDIENSEDNKSTLSLQFVDSAADMPLAKMHGAFSTNVVASKELQQPGSARSTRHLEIELPKEASYQEGDHLGVIPRNYEGIVNRVTARFGLDASQQIRLEAEEEKLAHLPLAKTVSVEELLQYVELQDPVTRTQLRAMAAKTVCPPHKVELEALLEKQAYKEQVLAKRLTMLELLEKYPACEMKFSEFIALLPSIRPRYYSISSSPRVDEKQASITVSVVSGEAWSGYGEYKGIASNYLAELQEGDTITCFISTPQSEFTLPKDPETPLIMVGPGTGVAPFRGFVQARKQLKEQGQSLGEAHLYFGCRSPHEDYLYQEELENAQSEGIITLHTAFSRMPNQPKTYVQHVMEQDGKKLIELLDQGAHFYICGDGSQMAPAVEATLMKSYADVHQVSEADARLWLQQLEEKGRYAKDVWAG

*Shimazuella soli* P450 (*So*P450)

HHHHHHHHHHHHAGAGAGENLYFQGMDEANIIPQPKTYGPLGNIPLIDKDKPILSFMKLAEEYGPIFRLQTPGDSTIVVSGHELVKEVCDESRFLKSAEGPLEKVRAFGGDGLFTSWTHEPNWRKAHNILMPTFSQRAMKDYHDMMVDIAVQLIQKWIRLNPDETVDVPDDMTRLTLDTIGLCGFNYRFNSYYRETPHPFITSMVRALDEAMHQTQRLDLQDKLMIRTKRQFQHDIQVMFSLVDSIIAERRAGKNQKENDLLSRMLNVSDPETGEKLDDENIRYQIITFLIAGHETTSGLLSFALYFLLKNPDKLKKAYEEVDQVLTGSTPTYKQVLHLKYVRMILNEALRLWPTAPAFSLYAKEDTIIGGKYPVKKEQERITVLIPQLHRDKEAWGEDVEEFRPERFEDQNKVPHHAYKPFGNGQRACIGMQFALHEATLVLGMLLRHFEFIDYKDYQLDIKQTLTLKPGDFNIQVQPRNQPAFQQSVSVTEDVAAESKMEMKQVKDHFDQKSTIQGLNNRPLLVLYGSDTGTAEGIARELADTATLHGVHTEVATLNERIGELPKEGAVLIVTSSYNGKPPSNAGQFVQWLEAVQAGELSGVQYAVFGCGDHNWASTYQDVPRFIDKQLAEKGAVRFSARGEGDVSGDFEEQFDQWKEKMWSDAIEAFGLEISDDVKKDQNTLSLQFVKGTGGSPLARSYEAVYAKVVENRELQSPDSGRSTRHIEITLPKGVSYQEGDHLGVLPANSKENVHRVLQKYKLNENDQVVLTASGRSMAHLPLAQEVSLRDLLLYSVELQDAATRAQIRELAAFTVCPPHKRELEALLEEGIYQEQVLKKRISMLDLLEKYEACEMPFERFLELLHPLKPRYYSISSSPRLNSERASITVAVVRGPAWSGLGEYRGVASNYLADCKPGEDVMMFIRTPESNFQLPEESETPIIMVGPGTGLAPFRGFLQARDAMKQEGKTLGEAYLYFGCRNEADFIYRKELEQYKKNGIMTLYIAFSRKEGIPKTYVQHVMASNAESLIRILDQGGRLYICGDGSRMAPEVEATLKNSYQEVHGAGEQEASQWLEKLQKDGQYAKDVWAGL

*Thermothelomyces thermophilus* ATCC 42464 P450

HHHHHHENLYFQGAGAGAGAGAGMADKTTETVPIPGPPGLPLVGNALAFDSELPLRTFQEFAEEYGEIYRLTLPTGTTLVVSSQALVHELCDDKRFKKPVAAALAEVRNGVNDGLFTAREEEPNWGIAHRILMPAFGPASIQGMFTEMHEIASQLALKWARHGPDTPIFVTDDFTRLTLDTLALCTMNFRFNSYYHDELHPFINAMGNFLTESGARAMRPAITSIFHQAANRKYWEDIEVLRKTAQGVLDTRRKHPTNRKDLLSAMLDGVDAKTGQKLSDSSIIDNLITFLIAGHETTSGLLSFAFYLLIKHQDAYRKAQEEVDRVIGKGPIKVEHIKKLPYIAAVLRETLRLCPTIPIINRAAKQDEVIGGKYAVAKDQRLALLLAQSHLDPAVYGETAKQFIPERMLDENFERLNREYPDCWKPFGTGMRACIGRPFAWQEAVLVMAMLLQNFDFVLHDPYYELHYKQTLTTKPKDFYMRAILRDGLTATELEHRLAGNAASVARSGGGGGGPSKPTAQKTSPAEAKPMSIFYGSNTGTCESLAQRLATDAASHGYAAAAVEPLDTATEKLPTDRPVVIITASFEGQPPDNAAKFCGWLKNLEGDELKNVSYAVFGCGHHDWSQTFHRIPKLVHQTMKAHGASPICDEGLTDVAEGNMFTDFEQWEDDVFWPAVRARYGAAGAVAETEDAPGSDGLNIHFSSPRSSTLRQDVREATVVGEALLTAPDAPPKKHIEVQLPDGATYKVGDYLAVLPVNSKESIGRVMRKFQLSWDSHVTIASDRWTALPTGTPVPAYDVLGSYVELSQPATKRGILRLADAAEDEATKAELQKLAGDLYTSEISLKRASVLDLLDRFPSISLPFGTFLSLLPPIRPRQYSISSSPLNDPSRATLTYSLLDSPSLANPSRRFVGVATSYLSSLVRGDKLLVSVRPTHTAFRLPDEDKMGETAIICVGAGSGLAPFRGFIQERAALLAKGTQLAAALLFYGCRSPEKDDLYRDEFDKWQESGAVDVRRAFSRVDSDDTEARGCRHVQDRLWHDREEVKALWDRGARVYVCGSRQVGEGVKTAMGRIVLGEEDAEDAISKWYETVRNDRYATDVFD

*Marinactinospora thermotolerans* P450

HHHHHHENLYFQGAGAGAGAGAGMTVTEIPAPRGLPFVGNTFQIPAHAPSAYFNSLAAAHPEGIYRVTVLGQEMLLVYDPDLVAEVCDESRFYKSIDPPLSIVRDFGGDGLFTARRSEPVWGHAHRILMPAFGQRSMKAYFPQMLEIADQLVASWRRRQGEEIDVSDDMTRLTLDTISLTGFDYRFDSFDSPELHPFLQAMGRALTEAMLRSRRPPFVTRFMKRQERGYRADIATMRELVDDIIRHRRQEGRAGGRDLLGLMMEAADPKTGARLSDENIRNQVLTFLIAGHETTSGLLSFAVYNLLRNPHVLARAYAEVDRLLPGEEPPTYEAIMRLDVIPRILDETLRIWSPIPAFSVKSFVDTTLGGHPVPKGRKVVVLLPSLHRHPKAWDRPEEFDIDRWLPENRAGHHPAGYKPFGNGQRACIGRQFALTEARLALALVLRSFAISDPHAYRMRTRQTLTLKPEGFTIRVRERRPHERAAVVPAEPEPETGGDEVTVTGVALTVAYGSNLGTAADVAERLAERAGRSGFTTTLTTLDELAAAPPGEGVLAVVTSTYNGKAPDNAQDFDALAALPSMKGVRLALLGCGNTQWPTYQDFPRRAFDKLVRAGAEPLLERCEADTDGDFDGAVSEWTTRLWAALAAHYGTSAAQEGPRYTMEVLPETEVRPAVVSANAVPLTVVANEELTGDPEGLWDFELEAPRPGVRSIVAELPEGTAYTAGDHLAVFAKNDPTAVERALRLLRIPREQVVRLRAEGSSHLPLDTPVTAGLLLSEFVELQETATRADLETLAAHTACPWTRGELARLDYVEDVLTPRVSVLTLLERFPAIELPLPVFLEMAGPIRPRYYSISSSALADPGRVRITVGLVEGPALSGAGHYRGTCSAYLAGLRPGDVFYGYVRVPAPAFRLPADPATPLLLVGPGTGFAPLRGFLEERFLTGASGRVEVFCGCRHPEHDWLYRRELEEWERAGVARVHTAFSAVEGHPHRFVQDALAAKADVVGDLLDQGAHVYVCGDGVRMAPAVRRTLARIHHDRTGGDGEAWMRRLEAEGRYHQDIFA

*Thermocatellispora tengchongensis* P450

HHHHHHHHHHHHAGAGAGENLYFQGMLQVPAHSPSAYFNELAGRHPQGIYRLDLMGRDILFVYDPDLVAELCDESRFYKSIDPPLSIVRDFAGDGLFTARHDEPVWGQAHRILMPAFSQRSMKAYFPQMLEIAQNLVASWRRRQGQDLNVPDDMTRLTLDTISLTGFDYRFDSFESAELHPFLRAMGRALTEAMLRNRQLPLVTKLKRRREEGYRADIALMRELVDDVIRQRRAAGKAGTGDLLGLMLQAADPRTGERLSDANIRDQVLTFLIAGHETTSGLLSFALYNLLRNPHALALAYAEVDRLLPGDEPPTYETIMKLDVIPRVLEETLRLWSPIPSFAVTAYADTTLGGHPIAAEQRCVILLPPLHRHPKAWDRPEEFDIDRWLPERKREHHPAAYKPFGNGERACIGRQFALTEARLALAMVLREFALADPGAYRMKIKQTLTLKPEDFTIRVRARQPHERAVPAAQGEAAEDAPRDLAAEVTVTGVSLAVAYGSNLGTSADIAERLAERARHAGFATTLLTLDELAAAPPREGLLAVVTSTYNGKAPDNAQEFDALEELPRMDGVRLALLGCGNTQWPTYQGFPKRAFAKLTAAGAVPLVERGEADADGDFDGAVSAWTSALWAALAAEYGAGAAPAGPRYEMEVLTESEIRPAVVSERAVPLTVVSNEELTGDPTGLWDFSLEAPRPGVRSIVAELPEGVTYEPGDHLAVFARNEPELVERALRALRVPRDQVVRLRAHGATHLPVDTPVTAGLLLADFAELQDVATRADLATLAAHTRCPWTRGELDRLTAAYPDEVLAKRVSVIDLLERYPAIALPLPVFLEAAGPIRPRYYSISSSPLAAPRQVRITVGLVEGPAWSGADAAYRGMCSAYLAGLAPGEVFYGYVRVPAPPFRLPEDPATPIVLVGPGTGFAPLRAFLEHRALSAPGAEPAEVFTGCRHPGHDRLYHADLDAWERAGVARVHTAFSALPGHPHRYVQDALSANADTVWALLERGAHVYVCGDGLHMAPAVRRTLAAIYAERTGRDGDAWLRELEEAGRYQQDVFA

*Streptomyces cattleya* P450

HHHHHHHHHHHHENLYFQGAGAGAGAGAGMSPTPHSASGTTGAAAATPGAASPAPPVPVADISDTGFGTTPIQQAMALAREHGPVFRRRFGTFESLLVGSVDAVTELCDDERFVKAVGPVLTNVRQIAGDGLFTAYNDEPNWAKAHDILLPAFALSSMHTYHPTMLRVAKRLIAAWDTALADGAPVDVADDMTRMTLDTIGLAGFGYDFGSFRRGEPHPFVAAMVRGLLHSQALLSRKADDGVDHSAADEAFRADNAYLAQVVDEVIEARRASGETGTDDLLGLMLGAPHPSDGTPLDAANIRNQVITFLIAGHETTSGALSFALYYLAKNPAVLRRAQAEVDALWGDDPDPEPDYTDVGRLTYVRQVLNEALRLWPTAAAFGRQAVTDTVLDGRVPMRAGDTALVLTPVLHRDPVWGDNVEAFDPERFSPEREAARPVHAFKPFGTGERACIGRQFALHEAVMLLGMLIHRYRFLDHADYRLRVRETLTLKPDGFTLKLARRTSADRVRTVASRAAEGTAGQDAGLPTTARPGTTLTVLHGSNLGACREFAAGLADLGERCGFETTVAPLDAYRAGDLPRTSPVVVVAASYNGRPTDDAAGFVSWLEQAGPGAADGVRYAVLGVGDRNWAATYQKVPTLIDERLAECGATRLLERAAADAAGDLAGTVRGFGEALRRALLAEYGDPDSVGAVAGAEDGYEVTEVTGGPLDALAARHEVVAMTVTETGDLADLTHPLGRSKRFVRLALPDGATYRTGDHLAVLPANDPALVERAARLLGADPDTVLGVRARRPGRGTLPVDRPVTVRELLTYHLELSDPATAAQIAVLADRNPCPPEQAELKKLAPGRASVLDLVERYPALTGRLDWPTVLGTLLPQIRIRHYSVSSSPAVSPGHVDLMVSLLEADGRRGTGSGHLHRVRPGDVVYARVAPCREAFRIAAGDEVPVVMVAAGTGLAPFRGAVADRVALRSAGRELAPALLYFGCDHPEVDFLHAAELRGAEAAGAVSLRPAFSAAPDGDVRFVQHRIAAEADEVWSLLKGGARVYVCGDGSRMAPGVREAFTALYASRTGATAEQAAGWLADLVARGRYVEDVYAAG

*Aspergillus thermomutatus* P450

HHHHHHENLYFQGAGAGAGAGAGMSESQTIPIPGPRGVPLLGNIYDIEQEVPLKSIDLLADQYGPIYRLTTFGRSRVFISTHELVDEVCNEERFTKVVSAGLNEIRNGVHDGLFTANYPGEENWAIAHRVLVPAFGPLSIRGMFDEMYDIATQLVMKWARQGPTAPIMVTDDFTRLTLDTIALCAMGTRFNSFYHEEMHPFVEAMVGLLQGSGDRARRPALINNLPTGENAKYWNDITFLRNLAQELVETRRKNPEDKKDLLNALILGRDPKTGQGLTDDSIIDNMITFLIAGHETTSGLLSFLFYYLLKNPRAYKKAQEEVDSVIGRRKITVEDMSKLPYINAVMRETLRLRSTAPLIAVHAHPEKNKEDPVTLGNGKYVLNKDEAIVIILDKLHRDPQVYGPDAEEFKPERMLDENFEKLPKNAWKPFGNGMRACIGRPFAWQEALLVVAILLQNFNFQMDNPSYDLRIKQTLTIKPKDFHMRAALRHGLDATKLGIVLSGSADSAPPESSGAASKGRKQAAPRPGQLKPMHIFFGSNTGTCETFARRLADDAVGYGFAAEVQSLDSPMQNVPRDEPVVFITASYEGQPPDNAAHFFEWLSALKGNELEGVNYAVFGCGHHDWQATFHRIPKAVNQLVAEHGGNRLCDIGLADAANSDMFTDFDSWGESAFWPAITAKFGGGKSDEPQSSSSLQVEVSSGMRASTLGLQLQEGFVVENQLLSKPGVPAKRMIRFKLPSDMSYQCGDYLAVLPVNPSSVVRRAIRRFDLPWDAMLTIRKPTQAPKGSTAIPLDTPISAFELLSTYVELSQPASKRDLNALADAAVTDADVQAELRYLASSPTRFTEEIVKKRMSPLDILIRYPSIKLPVGDFLAMLPPMRVRQYSISSSPLADPSECSITFSVLNAPSLAAVSLPPAERAEAEQYMGVASTYLSELKPGERAHITVRPSHSGFKPPVDLKTPMIMACAGSGLAPFRGFVMDRAEKIRGRRSSVGSDAQLPEVGQPARALLYVGCRTQGKDDIHAAELAEWAQLGAVDVRWAYSRPEDGSKGRHVQDLMLEDREELVSLFDQGARIYVCGSTNVGNGVRQACKDIYLERRRQLRREARERGEDVAAEDDEDAAAEKFLDNLKTKERYATDVFT

*Pseudonocardia thermophila* P450

HHHHHHENLYFQGAGAGAGAGAGMTLALDDVPGPRGLPVLGNVFDVDTQDPIGGLVQLAEQYGPVYRLSLPQGSRLIVSSAELVAEVCDDERWDKHVGGGLSNIGGSGAGLFTAETGDPLWARAHNILMSPFSLPSMRDYLPKMIDLAEQLATKWERVNPGEEVDVPADMTRLTLDTIALCGFGYRFNSFYRETPHPFVDAMVRTLLEAQTRARQPKIAARLRIRAQRQAEEDLAFMNGLVDGLIAERRAQGDTADTSDLLGRMLTGVDSRTGERLPDDNIRAQCITFLIAGHETTSGLLSFAINYLLKEPAATERAKAEVDEVLGDRAEPTFEQIQRMTYVRQVLDESLRLWPTAPAFTRHPLADTVLGGYRVPAGTPVSVLIPALHRDPAAWGPDADRFDPDHMAAERIAALPPHAYKPFGTGQRACIGRQFALQEAVLVLATLLQRFELIDSRDYQLRTKMTLTVKPEDFHIQVRPRGRTLRRAEVPAPAAASAPAPTPEVARHGTPLAVLFGSNLGTAEAIATTLAAEGGDRGFAVTLGALDDHVGDLPEGGAAIVVSSSYNGTPPDNAAGFCRWICGDPAVPGTAYTVFGCGSTEWASTYQAVPILLDERLEATGGRRVHPRGAADARGDTDAAYRAWREGLWASLAAALELPAEVGERIDTGPRLAVTLANRQTTNPVVLSYRARPARIRSNVELMPSQNGKPPERSVRHLTIELPEDLTYRAGDHLGVLPRNGIDMIRRVIVRFGLDAGQYATIIPNGGVFTHLPVDEPAPLLGVLGSCVELQDTATRSDIEALARYAPPDVRPELEALAGDDYAVHVGEPNRSVLDLLEEFPTCALPFAEFLDMLPPLRPRYYSISSSPLVDAGTCTITAGVLRAPARSGVGTFTGVCSGHLAALPELGTAFVFVREPSIPFRPPENPHIPMIMIGAGTGLAPFRGFLQERAAQRAQGAPVSRSLLFAGCRGPAIDQLYGDELESWSDIVDVEYAYSRAGDRRRYVQDAIRDRAETVWGLLQQGAPVFVCGNAATMAPGVRAALTDVFRDRTGTGQADADAWLAGLRGSDRFLEDIWGG

*B. nakamurai* P450

HHHHHHENLYFQGAGAGAGAGAGMKQLNAIPQPKTYGPLKNLPHIEKEKLAQSLWKIAEEYGPIFRFEFPSSAAVFVSGHRLAAEVFDESRFDKNLGKGLLKVREFAGDGLFTSWTNEKNWQKAHRILLPSFSQKAMKGYHSMMLDIAMQLVQKWSRLNPKEEIDVAEDMTRLTLDTIGLCGFHFRFNSFYRDTQHPFIISMLRGLQEAMRQSQRHSLQDKLMVKTRHQFQQDIEVMNSLVDKIIAERRENPDEEITDLLSLMLDAKDPVTGETLDDENIRYQIITFLIAGHETTSGLLSFAIYCLLKNKDKLEKASQEAEQVLTGDLPDYKQIQQLKYIRMVLNETLRLYPTAPAFSLYAKEDTVLGGKYPIEKGQPVTILTPQLHRDKSAWGEDAEMFRPERFSDPAAIPNHAYKPFGNGQRACIGMQFALHEATMVLGLVLKHFELIDHKHYELSIKEALTIKPGDFTIRVKKKDVPLQPVQEKQAETSDTKEELKDIPTHGTPLLVLYGSNLGTAEGIAEELFDIGRSKGFAAETAPLDDYIGKLPTEGAVVIVTASYNGAPPDNAAGFVKWMDTLKDHDLNGVSYAVFGCGNRNWASTYQRIPRLIDKTLELKGAKRLTSIGEGDGADDFESSQEAWENLFWQDIIKAFDLEDVTDQQNRSSLSIEFVNEATETPLAKTYDAFETKVIKNKELHTGNSTRSVRHIELLLPETEAYQEGDHLGVLPKNSDELINRVIRRFGLDPHQLFKISGRHVPHLPMDRPVNASELLASHVELQEPATRAQLRELAAYTVCPPHQKELEYLHSDDAAYRENVLKKRMTMLDLLEDYPACEMPFERFLELLPSLKARYYSISSSPKAANEKVSITVGVVAAPSWSGRGEYRGVASNYLAGLQEGDRAVCFIRSPQSGFALPENTETPLIMVGAGTGIAPFRGFIQARAAEKAAGNSLGEAHLYFGCRHPEEDDLYKDEFIQAEQSGLVTVHRAYSRLDQNCKIYVQDILHREASQIIGLLDKGGHLYICGDGSKMAPAVENVLLRAYESVHDTDANTSLKWLEKLQAEGRYAKDVWAGV

*Bacillus cereus* P450

HHHHHHHHHHENLYFQGAGAGAGAGAGMEKKVSAIPQPKTYGPLGNLPLIDKDKPTLSFIKIAEEYGPIFQIQTLSDTIIVVSGHELVAEVCDETRFDKSIEGALAKVRAFAGDGLFTSETHEPNWKKAHNILMPTFSQRAMKDYHAMMVDIAVQLVQKWARLNPNENVDVPEDMTRLTLDTIGLCGFNYRFNSFYRETPHPFITSMTRALDEAMHQLQRLDIEDKLMWRTKRQFQHDIQSMFSLVDNIIAERKSSGDQEENDLLSRMLNVPDPETGEKLDDENIRFQIITFLIAGHETTSGLLSFAIYFLLKNPDKLKKAYEEVDRVLTDPTPTYQQVMKLKYMRMILNESLRLWPTAPAFSLYAKEDTVIGGKYPIKKGEDRISVLIPQLHRDKDAWGDNVEEFQPERFEELDKVPHHAYKPFGNGQRACIGMQFALHEATLVMGMLLQHFELIDYQNYQLDVKQTLTLKPGDFKIRILPRKQTISHPTVLAPTEDKLKNDEIKQHVQKTPSIIGADNLSLLVLYGSDTGVAEGIARELADTASLEGVQTEVVALNDRIGSLPKEGAVLIVTSSYNGKPPSNAGQFVQWLEELKPDELKGVQYAVFGCGDHNWASTYQRIPRYIDEQMAQKGATRFSKRGEADASGDFEEQLEQWKQNMWSDAMKAFGLELNKNMEKERSTLSLQFVSRLGGSPLARTYEAVYASILENRELQSSSSDRSTRHIEVSLPEGATYKEGDHLGVLPVNSEKNINRILKRFGLNGKDQVILSASGRSINHIPLDSPVSLLALLSYSVEVQEAATRAQIREMVTFTACPPHKKELEALLEEGVYHEQILKKRISMLDLLEKYEACEIRFERFLELLPALKPRYYSISSSPLVAHNRLSITVGVVNAPAWSGEGTYEGVASNYLAQRHNKDEIICFIRTPQSNFELPKDPETPIIMVGPGTGIAPFRGFLQARRVQKQKGMNLGQAHLYFGCRHPEKDYLYRTELENDERDGLISLHTAFSRLEGHPKTYVQHLIKQDRINLISLLDNGAHLYICGDGSKMAPDVEDTLCQAYQEIHEVSEQEARNWLDRVQDEGRYGKDVWAGI

Green, His-tag; purple, tobacco etch virus protease cleavage site; red, (AG)*_n_* linker.

**Supplementary Table 2. Oligonucleotides used in this study**

| **Primer name** | **Sequence (5'🡪 3')** |
| --- | --- |
| P450BM3/A83F-F | GATTTTtttGGAGACGGGTTATTTACAAGC |
| P450BM3/A83F-R | GTCTCCaaaAAAATCACGTACAAATTTAAG |
| P450BM3/A83F-CR^I^-27-F | CCTTCAAGCACTGAACAGTCTGC |
| P450BM3/A83F-CR^I^-27-R | GTTCAGTGCTTGAAGGAATACCGCC |
| P450BM3/A83F-CR^I^-26-F | AGCAGACTGTTCAGTTGAAGGAATACCGCCAAGCGGAATTTTTTT |
| P450BM3/A83F-CR^I^-26-R | ACTGAACAGTCTGCTAAAAAAGTACGCAAAAAGGCAGAAAAC |
| P450BM3/A83F-CR^I^-25-F | TTTAGCAGACTGTTCTGAAGGAATACCGCCAAGCGGAATTTT |
| P450BM3/A83F-CR^I^-25-R | GAACAGTCTGCTAAAAAAGTACGCAAAAAGGCAGAAAACGCT |
| P450BM3/A83F-CR^I^-24-F | CAGTCTGCTAAAAAAGTACGCAAAAAGGCAGAAAACGCTCATAAT |
| P450BM3/A83F-CR^I^-24-R | TTTTTTAGCAGACTGTGAAGGAATACCGCCAAGCGGAATTTT |
| P450BM3/A83F-CR^I^-23-F | TCTGCTAAAAAAGTACGCAAAAAGGCAGAAAACGCTCATAAT |
| P450BM3/A83F-CR^I^-23-R | TACTTTTTTAGCAGATGAAGGAATACCGCCAAGCGGAATTTT |
| *So*P450/G84F-F | TGCATTTTTCGGTGATGGCCTGTTTACCAGCTGGACCCAT |
| *So*P450/G84F-R | CCATCACCGAAAAATGCACGCACTTTTTCCAGCGGACCTT |
| *So*P450/G84F-CR^I^-28-F | tttgatcagaaaagcaccattcagggcctgaataat |
| *So*P450/G84F-CR^I^-28-R | gcttttctgatcaaaggcaacatcttcggtcacgct |
| *So*P450/G84F-CR^I^-27-F | gcttttctgatcaaaaacatcttcggtcacgctcacgctctg |
| *So*P450/G84F-CR^I^-27-R | tttgatcagaaaagcaccattcagggcctgaataatcgtccg |
| *So*P450/G84F-CR^I^-26-F | gcttttctgatcaaaatcttcggtcacgctcacgctctgctg |
| *So*P450/G84F-CR^I^-26-R | tttgatcagaaaagcaccattcagggcctgaataatcgtccg |
| *So*P450/G84F-CR^I^-25-F | gcttttctgatcaaattcggtcacgctcacgctctgctgaaa |
| *So*P450/G84F-CR^I^-25-R | tttgatcagaaaagcaccattcagggcctgaataatcgtccg |
| *So*P450/G84F-CR^I^-24-F | gcttttctgatcaaaggtcacgctcacgctctgctgaaaggc |
| *So*P450/G84F-CR^I^-24-R | tttgatcagaaaagcaccattcagggcctgaataatcgtccg |
| *So*P450/G84F-CR^I^-23-F | gcttttctgatcaaacacgctcacgctctgctgaaaggccggc |
| *So*P450/G84F-CR^I^-23-R | tttgatcagaaaagcaccattcagggcctgaataatcgtccg |
| *So*P450/G84F-CR^I^-22-F | gcttttctgatcaaagctcacgctctgctgaaaggccggctg |
| *So*P450/G84F-CR^I^-22-R | tttgatcagaaaagcaccattcagggcctgaataatcgtccg |
| *So*P450/G84F-CR^I^-10-F | cagggcctgaataatcgtccgctgctggtgctgtatggt |
| *So*P450/G84F-CR^I^-10-R | attattcaggccctgaaaggccggctgattgcgcggctg |
| *So*P450/G84F-S593A-F | CAGCTACCTATCAGGATGTGCCGCGTTTTATTGATAAAC |
| *So*P450/G84F-S593A-R | TAGGTAGCTGCCCAATTATGATCGCCACAGCCAAAAAC |
| *So*P450/G84F-Q401A-F | TAATGGTGCTCGTGCATGCATTGGTATGCAG |
| *So*P450/G84F-Q401A-R | AGCACCATTACCAAACGGTTTATAGGCATGATGCG |
| *So*P450/G84F-R99A-F | AATTGGGCCAAAGCCCATAATATTCTGATGCCGACCTTTAG |
| *So*P450/G84F-R99A-R | GGCTTTGGCCCAATTCGGTTCATGGGTCCAGCTGGTAAAC |
| *So*P450/G84F-H102A-F | attggcgcaaagccgcaaatattctg |
| *So*P450/G84F-H102A-R | cagaatatttgcggctttgcgccaattcggttcat |

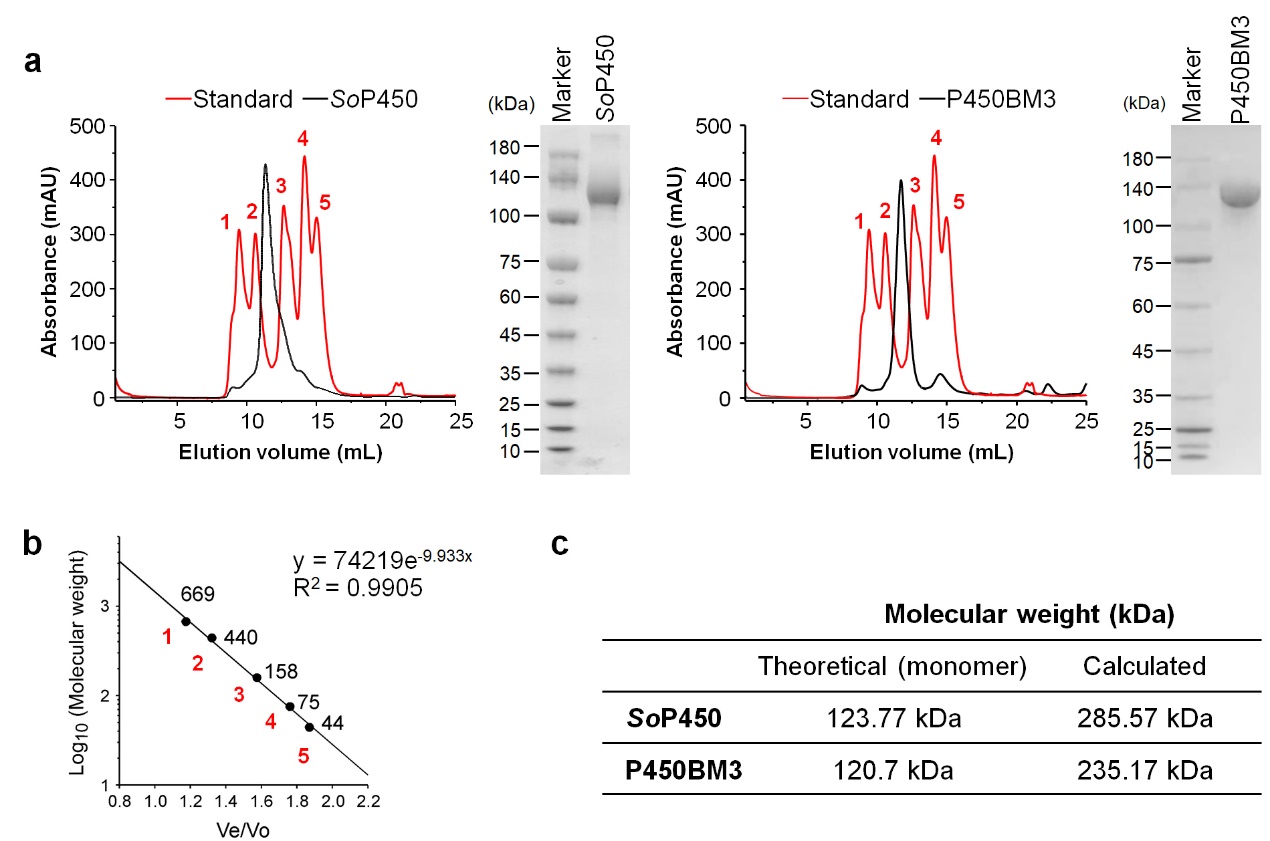

**Supplementary Fig. 1 |** Oligomerization status of *So*P450 and P450BM3 in the solution. **a**, The size-exclusion chromatographic analyses of *So*P450 and P450BM3. **b**, Molecular weight calibration curve for protein standards of thyroglobulin (1), ferritin (2), aldolase (3), conalbumin (4) and ovalbumin (5). **c**, The theoretical and calculated molecular weight of *So*P450 and P450BM3.

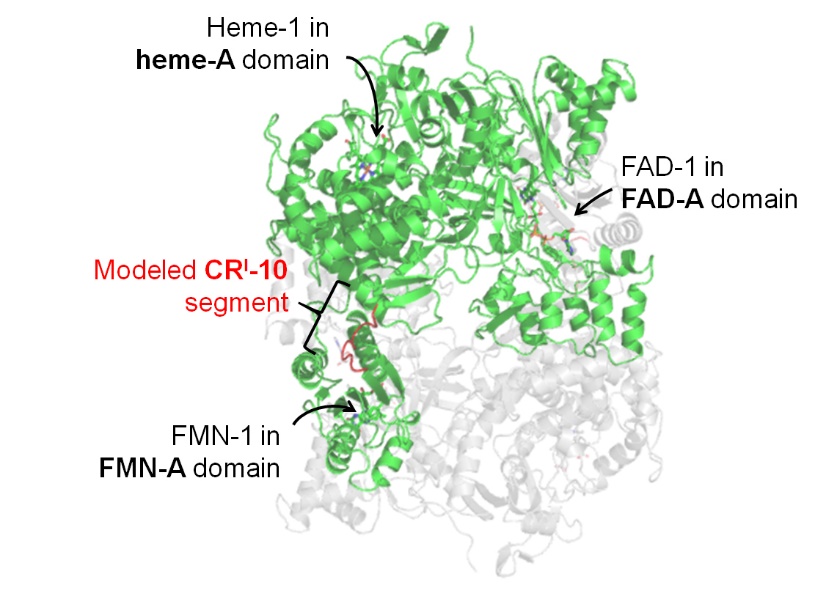

**Supplementary Fig. 2 |** The model of *So*P450-CR^I^-10. Chain A of *So*P450 homodimer is highlighted by green and chain B in gray, with the prosthetic groups (sticks) bound in three cognate domains indicated by arrows. The modeled CR^I^ segment is colored in red. The model construction processes are described in Supplementary Computation Information and **Supplementary Fig. 13**.

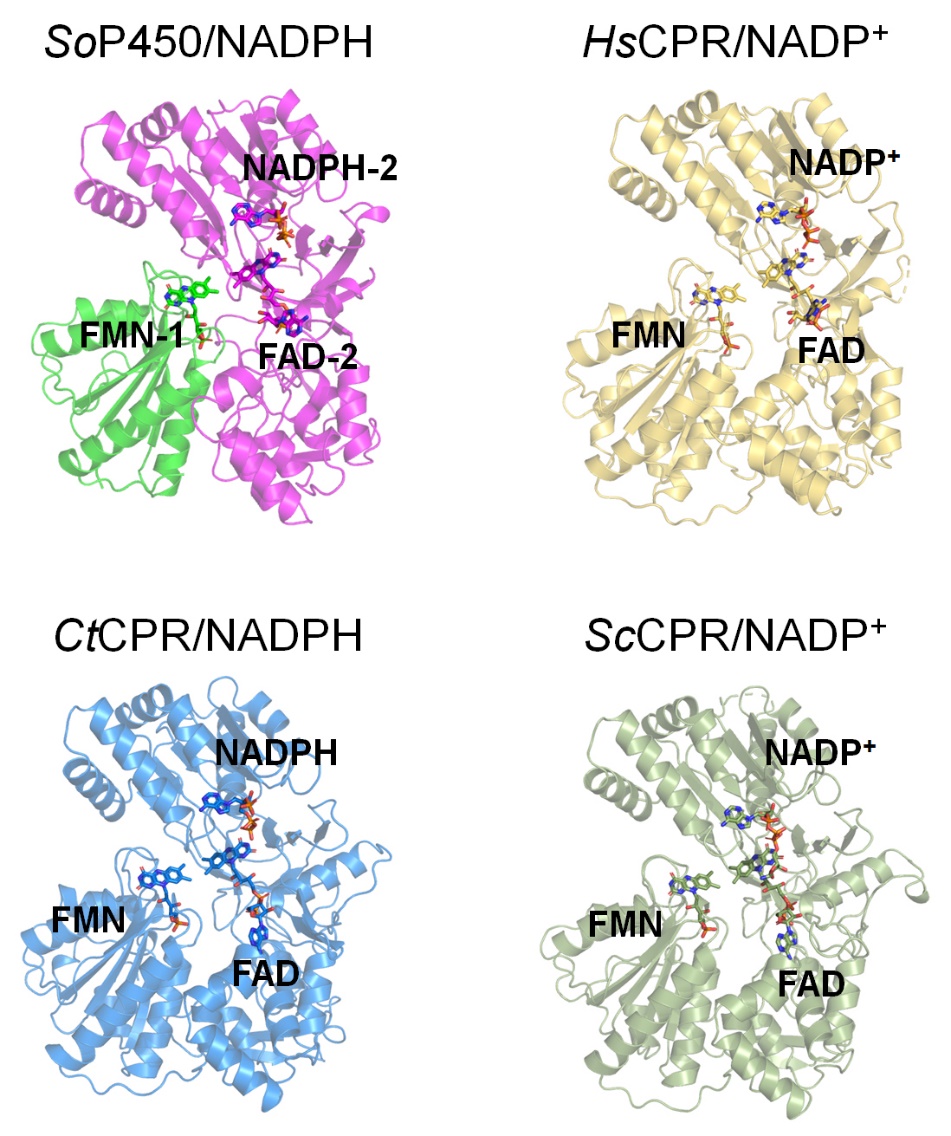

**Supplementary Fig. 3 |** The structures of CPR-domain of *So*P450 and NADPH:cytochrome P450 reductases (CPRs) from homo sapiens (PDB ID, 5FA6), *Candida tropicalis* (PDB ID, 6T1T) and *Saccharomyces cerevisiae* (PDB ID, 2BPO). The bound ligands are indicated and displayed as sticks. The FMN- and FAD-binding domain in *So*P450/NADPH from two monomers are colored differently.

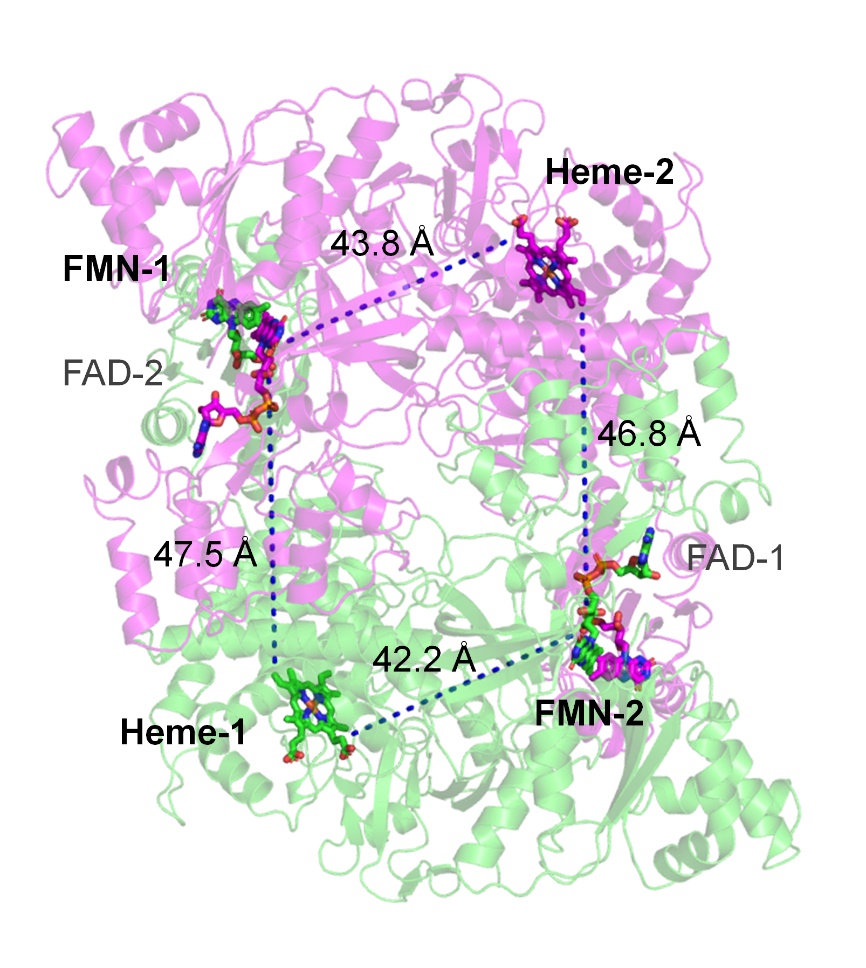

**Supplementary Fig. 4 |** The edge-to-edge straight-line distances between FMN and heme in the cryo-EM structure of *So*P450. Two polypeptide chains in the homodimer are colored in green and magenta, and the cofactors are displayed as sticks.

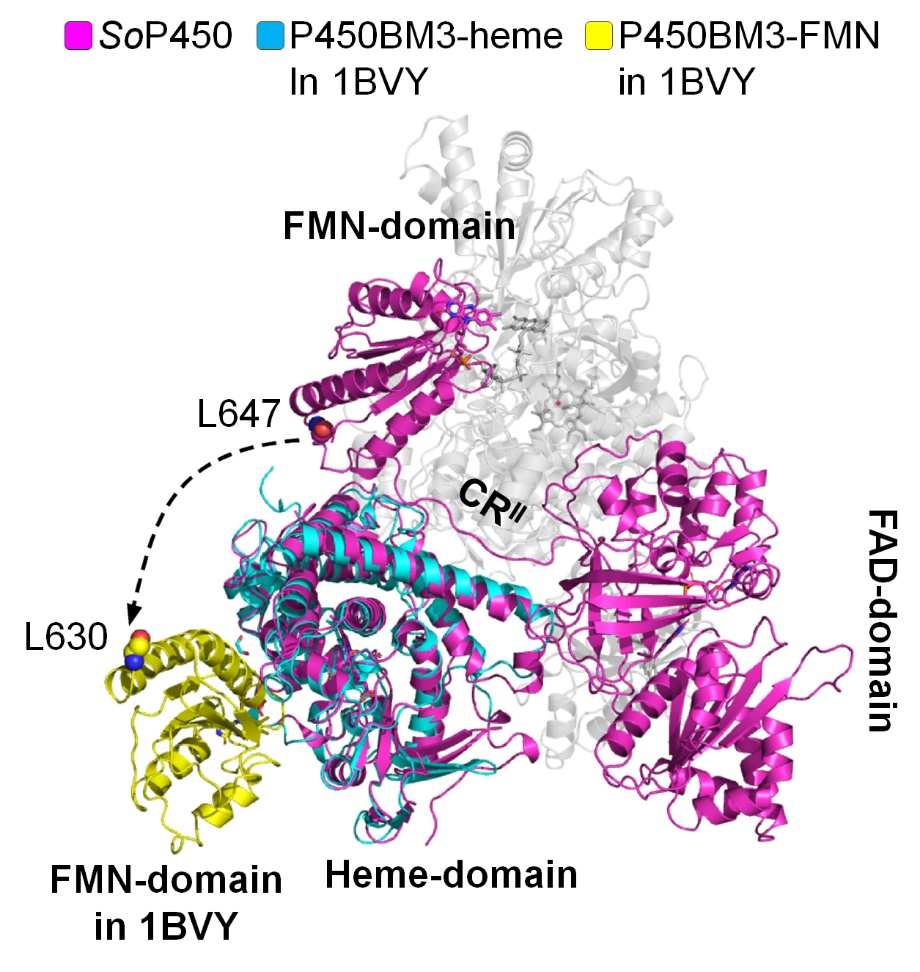

**Supplementary Fig. 5 |** Structural superimposition of cryo-EM structure of *So*P450 and the crystal structure of partial P450BM3 complex that contains FMN- and heme-domain (PDB ID, 1BVY). The main chain of L630, the residue at the most C-terminus of the FMN-domain of P450BM3 and that of the equivalent in *So*P450 (L647) are shown in spheres. The dash curve arrow depicts the trace if *So*P450 FMN-domain relocates to the position of P450BM3 FMN-domain in the complex structure.

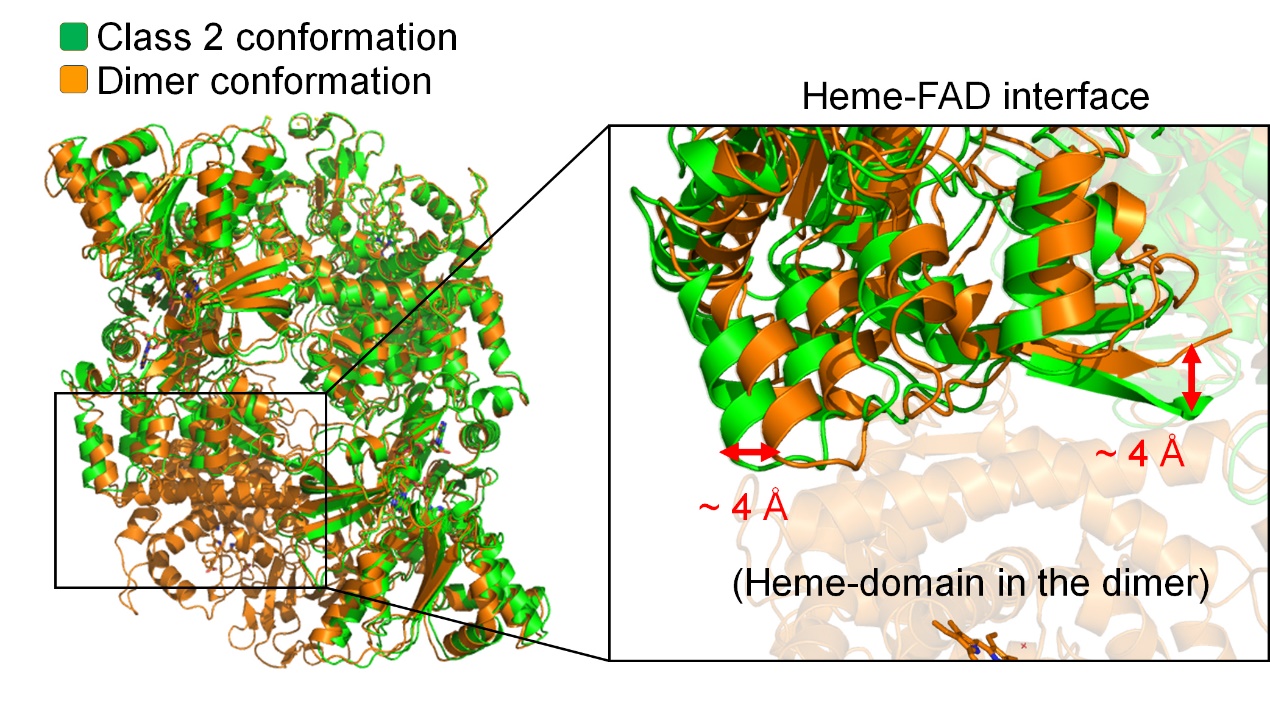

**Supplementary Fig. 6 |** Structure comparison of class 2 conformation and homodimeric form of *So*P450. The zoom-in view displays the interface between heme- and FAD-domain of two structures. The deviated distances in some displaced regions are noted by red arrows.

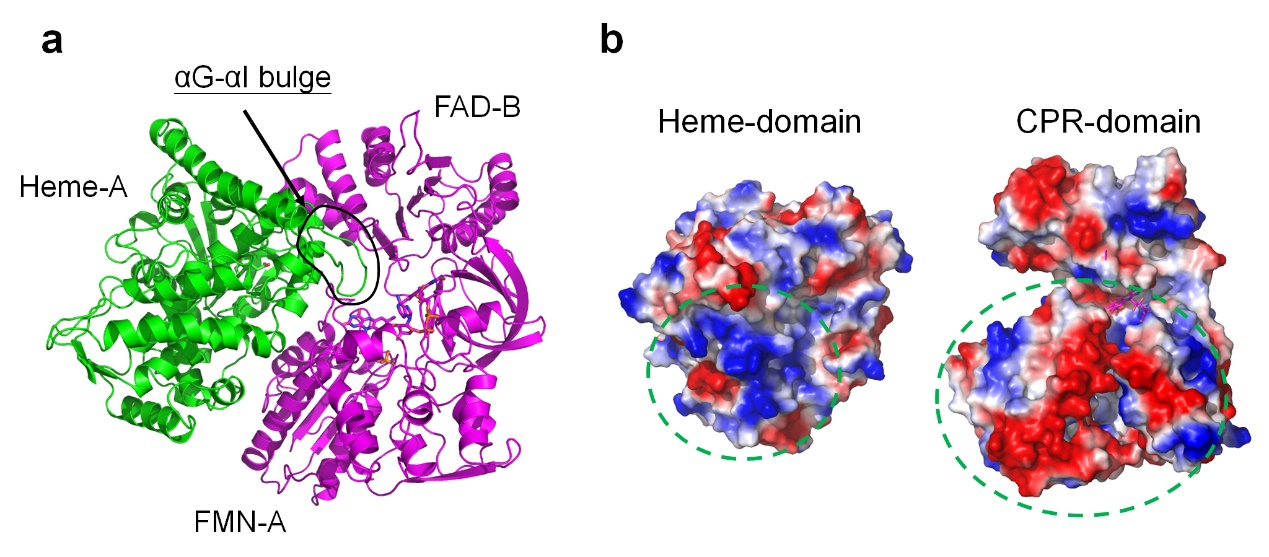
 **Supplementary Fig. 7 |** A partial model of *So*P450 in catalytic state displayed in **Fig. 5a**. **a,** The model of heme-A (green) and CPR-domain comprising FMN-A and FAD-B (magenta). The bulge formed between helix G and I (αG-αI bulge) is indicated. **b,** The electrostatic potential at the contact interface between two domains. The circled parts highlight the electrostatic complementary contact surfaces on the heme- and FMN-domain.

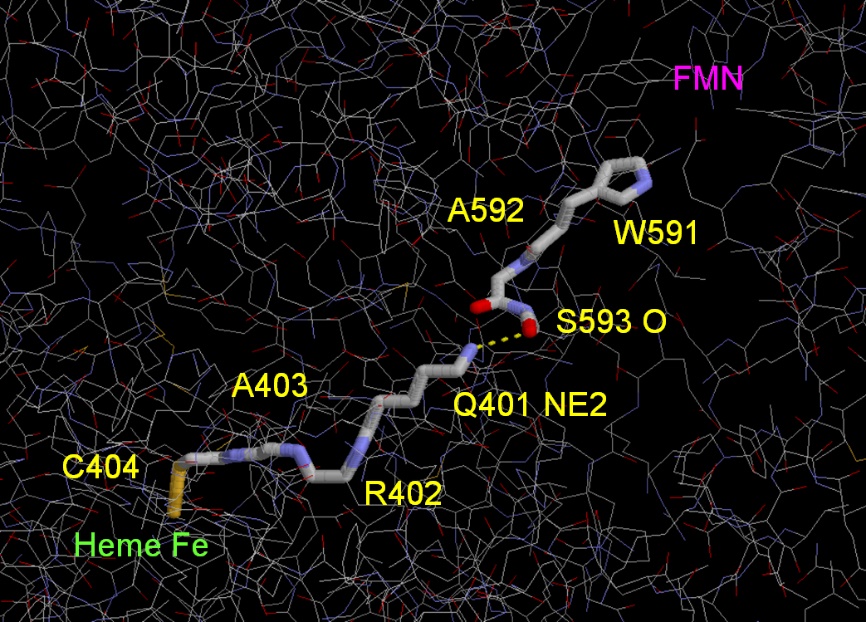

**Supplementary Fig. 8 |** **Electron traveling path predicted by program HARLEM.** The electron traveling path starting from the indole of W591 that stacks to FMN to the heme iron comprises A592, S593, Q401, R402, A403 and C404 revealed by program HARLEM based on the model of the catalytic status of *So*P450 displayed in **Fig. 5a**. Dash, the nonbonded jump between S593 O and Q401 NE2.

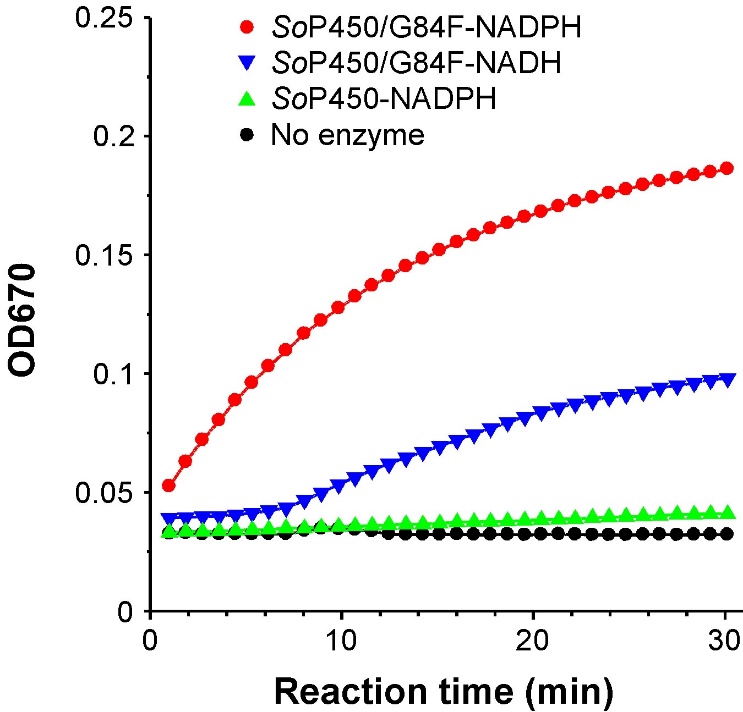

**Supplementary Fig. 9 |** Indigo production of wild type and variant G84F of *So*P450. 1 mL reaction containing 5 μM enzyme, 2 mM indole and 0.2 mM NADPH or NADH were incubated at 37 °C and indigo formation during 30 min was monitored through optical density at 670 nm. The representative data of a triplicate assay is presented.

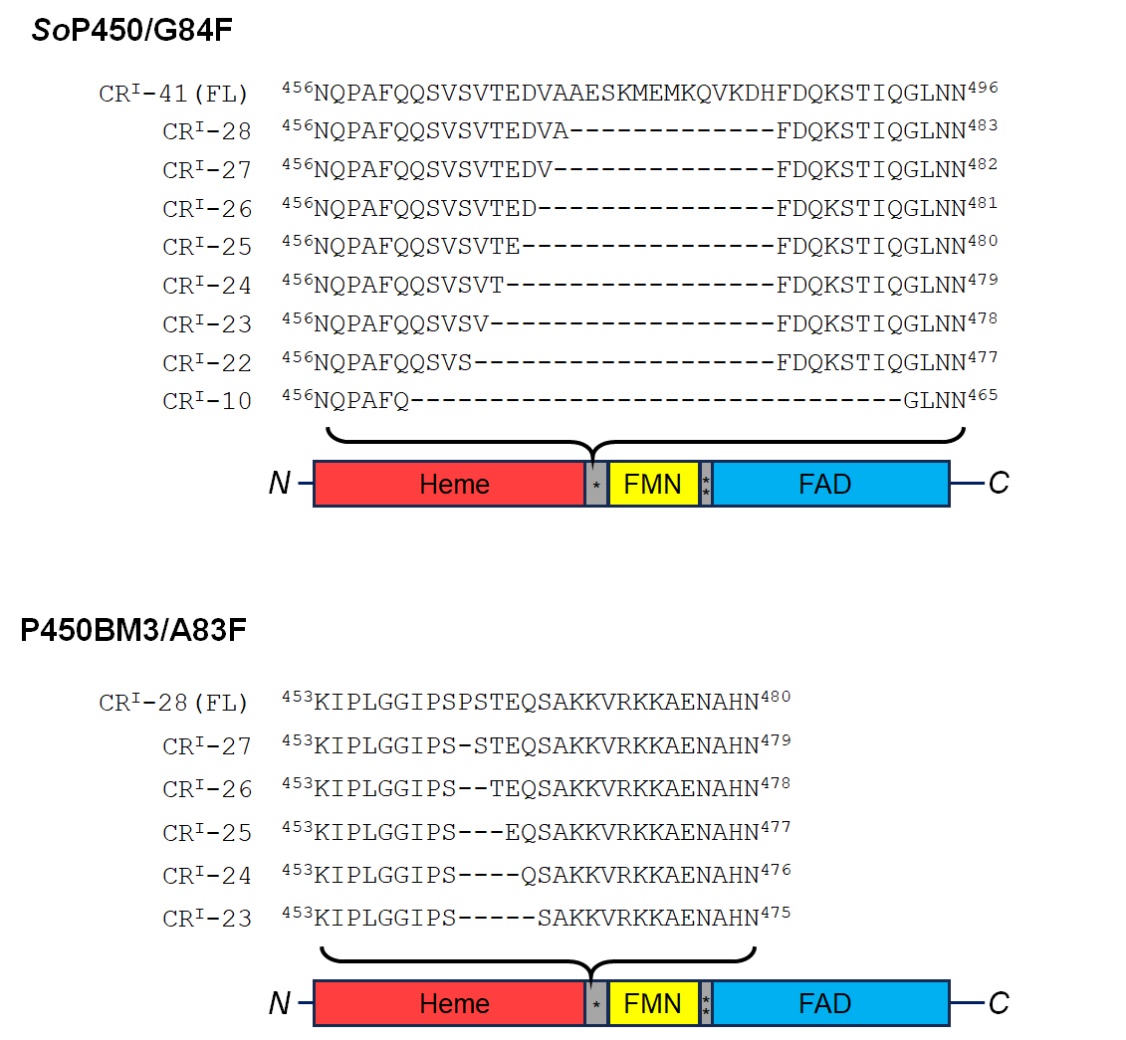

**Supplementary Fig. 10 |** Partial protein sequences of variant *So*P450/G84F and P450BM3/A83F that contain CR^I^ of varying length.

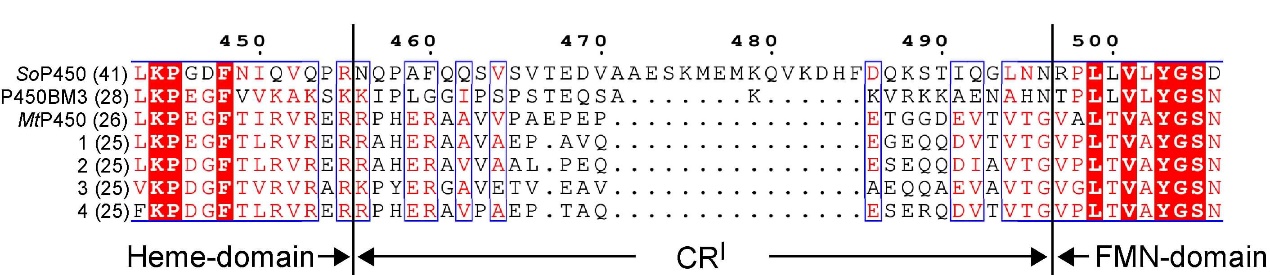

**Supplementary Fig. 11 |** Partial protein sequence alignment of *So*P450, P450BM3, *Mt*P450 and several CPR-containing self-sufficient P450s that possess short CR^I^ segment. 1, *Nonomuraea* sp. B12E4 (GenBank no., WP_347693761.1); 2, *Actinomadura parvosata* (GenBank no., WP_080040673.1); 3, *Streptosporangium* sp. NBC_01756 (GenBank no., WP_326825998.1); 4, *N. fastidiosa* (GenBank no., WP_378788029.1). The parenthesized numbers indicate the number of amino acids in the CR^I^ segment of each sequence. The sequences were aligned using Clustal Omega^1^ and the graphic was depicted using Espript 3.0^2^.

**Supplementary Computational Information**

1. The construction of *So*P450 models shown in **Extended Data Fig. 3**, in which CR^I^ is oriented via two different paths.

In the absence of homologous templates, the CR^I^ segment of *So*P450 was built *de novo* by Rosetta software as described in Methods. Two chains configured in heme-A linked FMN-A, or heme-B linked FMN-A, of the cryo-EM structure of *So*P450 were subjected to the computation. This procedure yielded a large ensemble of loop structures, illustrating that the reconstructed regions can adopt multiple conformations with considerable flexibility and ten representative models are displayed herein (**Supplementary Fig. 12**).

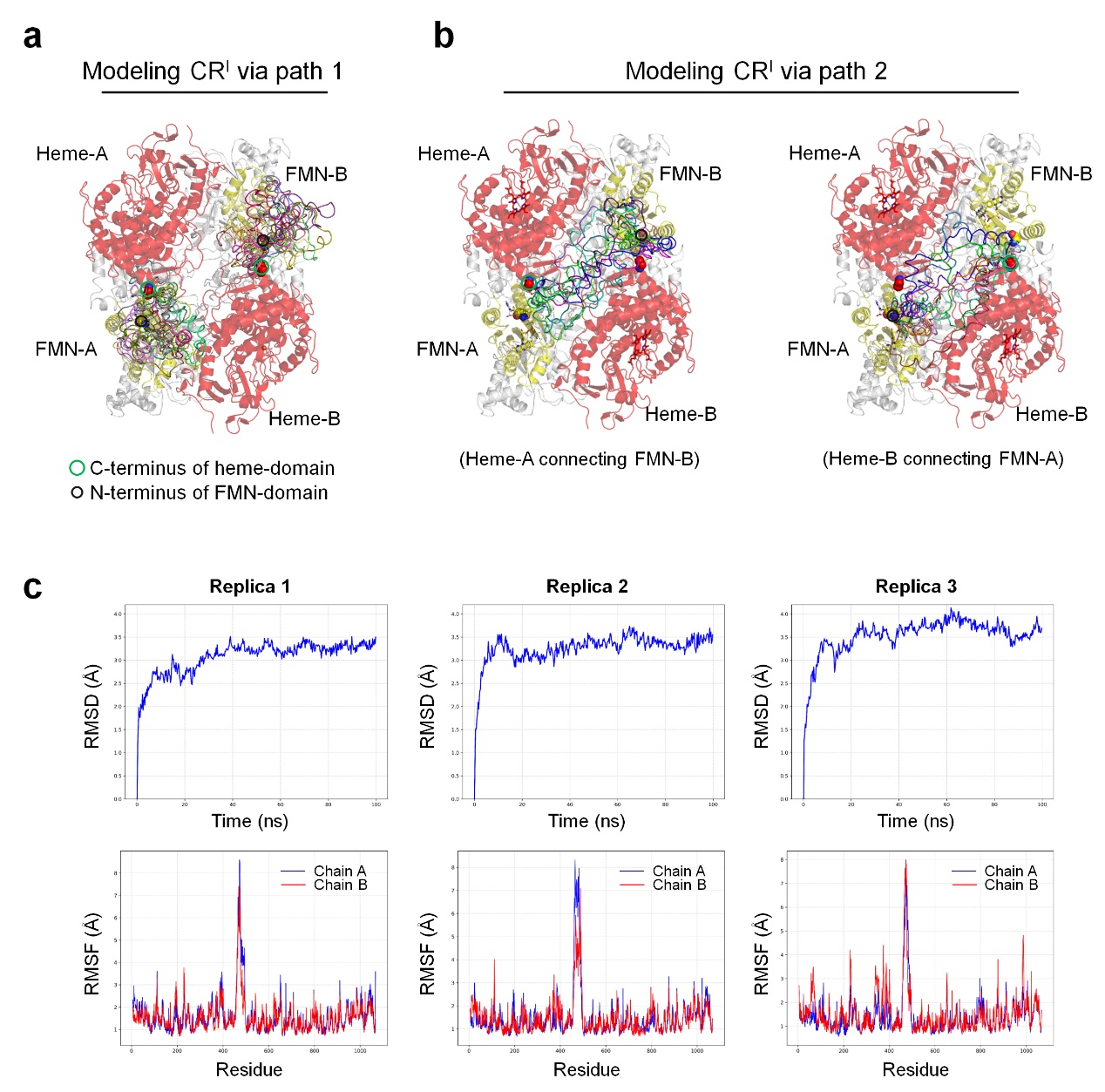

**Supplementary Fig. 12 |** The representative models of *So*P450 with heme- and FMN-domain linked via CR^I^ fragment oriented via **a**, path 1 and **b**, path 2. The structures of *So*P450 are displayed as described in **Extended Data Fig. 3**, with modeled CR^I^ fragments in various color. Note that the models of CR^I^ via path 2 that connect (**b**, left) heme-A and FMN-B and (**b**, right) heme-B and FMN-A are shown in two separate drawings to avoid confusion. **c**, The representative conformation of CR^I^ oriented via path 1 as shown in **Extended Data Fig. 3b** are selected for molecular dynamics (MD) simulations, during which all systems reached equilibrium (upper row). Analysis of the MD trajectories revealed pronounced fluctuations within the loop regions, as indicated by elevated RMSF values (lower row). Such dynamic behavior is consistent with experimental observations, where these flexible segments are often unresolved in the resolved structures.

2. The construction of *So*P450-CR^I^-10 models shown in **Supplementary Fig. 2**.

Remodeling of CRI segment that retains ten amino acids was performed based on the cryo-EM structure of *So*P450-CR^I^-10. The results indicated that a loop of this length was sufficient to connect heme-A and FMN-B (as well as heme-B and FMN-B) while clearly showed that a cross-link between heme-A and FMN-B (as well as heme-B and FMN-A) could not be formed.

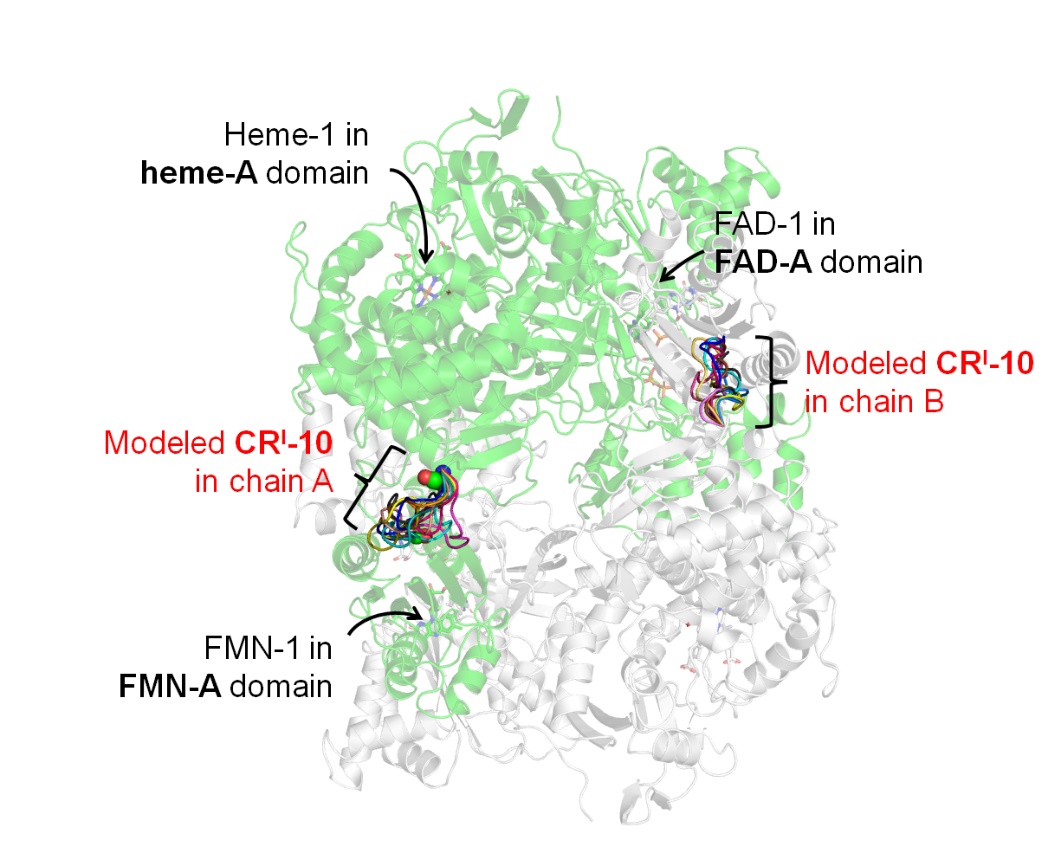

**Supplementary Fig. 13 |** The representative models of *So*P450-CR^I^-10 with heme- and FMN-domain linked via a ten-amino acid CR^I^ fragment oriented via path 1 illustrated in **Extended Data Fig. 3b**. The modeled CR^I^ fragments are in various color.
